# Neural network features distinguish chemosensory stimuli in *Caenorhabditis elegans*

**DOI:** 10.1101/2020.02.18.955245

**Authors:** Javier J. How, Saket Navlakha, Sreekanth H. Chalasani

## Abstract

Nervous systems extract and process information from their environment to alter animal behavior and physiology. Despite progress in understanding how different stimuli are represented by changes in neuronal activity, less is known about how they affect broader neural network properties. We developed a framework to use graph-theoretic features of neural network activity and predict ecologically-relevant stimulus properties – namely, stimulus identity and valence. Specifically, we used the transparent nematode, *Caenorhabditis elegans*, with its small nervous system, to define neural network features associated with various chemosensory stimuli. We trapped animals using a microfluidic device and exposed their noses to chemical stimuli known to be attractive or repellent, while monitoring changes in neural activity in more than 40 neurons in their heads. We found that repellents trigger higher average neural activity across the network, and that the tastant salt increases neural variability. In contrast, graph-theoretic features, which capture patterns of interactions between neurons, are better suited to decode stimulus identity than measures of neural activity. Furthermore, we show that a simple machine learning classifier trained using graph-theoretic features alone or in combination with neural activity features can accurately predict stimulus identity. These results indicate that graph theory reveals network characteristics that are distinct from neural activity, confirming its utility in extracting stimulus properties from neural population data.

**Significance Statement:** Changes in the external environment (stimuli) alter patterns of neural activity in animal nervous systems. A central challenge in computational neuroscience is to identify how stimulus properties alter interactions between neurons. We recorded neural activity data from *C. elegans* head neurons while the animal experienced various chemosensory stimuli. We then used a combination of activity statistics (i.e., average, standard deviation, and several frequency-based measures) and graph-theoretic features of network structure (e.g., modularity – the extent to which a network can be divided into independent clusters) to define neural properties that can accurately predict stimulus identity. Our method is general and can be used across species.

## Introduction

Animals have evolved mechanisms to encode the vast array of chemical information in their environment. The underlying neural circuitry encoding odor and taste information in both vertebrates and invertebrates is thought to include both labeled lines and combinatorial activity patterns. Specifically, odor information is initially filtered by olfactory sensory neurons that are organized into specific expression zones in the vertebrate olfactory epithelium [1], and into sensilla selective for pheromones [2], food odors [3], acids [4], oviposition cues [5] or toxic odors [6] in flies. This information is relayed to specific glomeruli and then higher-order centers in the brain [7, 8]. Similarly, taste information in both flies and mice are represented by spatial patterns of neural activity, likely using combinatorial coding [9–11]. While these studies highlight the progress made in understanding how chemical information is encoded in the periphery and early cortical areas, its processing and representation in higher brain centers is poorly understood. One solution to this problem is to monitor neural activity of the entire circuit in an intact nervous system as the animal experiences changes in its chemical environment, and to extract neural features that predict those changes.

The nematode *C. elegans*, with its nervous system consisting of just 302 neurons connected by identified chemical and electrical synapses [12, 13], is ideally suited to record neural activity across a large part of the network. *C. elegans* neurons express rapidly activating voltage-gated calcium channels, such that changes in neuronal calcium approximately correlate with neuronal depolarization [14–16]. By monitoring neural activity using genetically encoded calcium indicators, we previously showed that *C. elegans* sensory neurons encode chemical stimulus concentration and identity using a combinatorial code [17, 18]. Moreover, large-scale activity measurements in the *C. elegans* nervous system are aided by two innovations – custom-designed microfluidic devices that trap and precisely deliver chemical stimuli to adult animals during recording of neural activity [19, 20], and nuclear-localized genetically encoded calcium indicators that restrict fluorescent signals to easily resolved neuronal nuclei instead of overlapping cytoplasm [21].

Previous analyses of *C. elegans* whole-brain imaging data used principle components analysis (PCA) to show that neural activity lies in a low-dimensional space [22–24]. For example, Kato et al. [22] showed that the *C. elegans* neural network likely exists in a few global states that might represent locomotory commands, such as forward movement, reversals, turns, and others. Nichols et al. [23] and Skora et al. [25] revealed low-dimensional neural activity patterns associated with physiological and behavioral states, such as sleep and starvation. Critically, these approaches required having labels for neurons such that the same neuron can be uniquely identified across animals. However, in most model systems (e.g., recording from cortical neurons in a mouse), such neuron-specific labels do not exist, precluding the use of PCA-based approaches in this manner. Furthermore, Scholz et al. [26] found evidence that neural dynamics have higher dimensionality than previously thought, suggesting the engagement of many neurons in driving behavior. Hence, even with neuron-specific labels, it is important to probe how the entire network cooperates to process stimuli.

Therefore, here we ask: can we identify stimulus properties using graph-theoretic features of neural interactions recorded during stimulus onset or offset? Graphs are a natural representation to capture the pairwise interactions between nodes (or neurons) connected by edges (functional interactions). Graphs have been used to uncover structure-function relationships in physical, biological, social, and information systems [27–29]. By viewing a nervous system as a graph, we hypothesize that we may uncover more complex patterns of activity than if we considered neurons as independent units. Indeed, we identified a range of network features that emerge in response to chemosensory stimulation. We chose five chemical stimuli each at two different concentrations and monitored the responses of at least 40 head neurons. Head neurons in *C. elegans* include olfactory and gustatory sensory neurons, several downstream interneurons and command interneurons that direct locomotion, and some motor neurons [12]. We then computed how two stimulus properties – a chemical’s valence (attractant or repellent) and identity (i.e., chemical structure) – affect neural activity across the network. We observed that activity statistics and graph-theoretic features were successful in discriminating between stimulus properties, and validated these results using machine learning classifiers. Finally, we found that chemical identity mostly altered the subnetwork composed of putatively excitatory, as opposed to inhibitory, interactions, suggesting that patterns of excitation define the representation of a chemical in the nematode brain.

## Results

We performed whole-brain calcium imaging in 30 worms immobilized in an olfactory chip, a microfluidic device that permits near-instantaneous switching between two fluid flows (Fig. 1A; [19]). Each worm experienced three 21-minute imaging sessions: one without stimulation (“Spontaneous”, although M9 buffer is still present), one with buffer changes around the animal’s nose (“Buffer”, with M9 buffer), and one with chemical stimulation (“Stimulus”, where an odorant or tastant was diluted in M9 buffer). Sessions with stimulation had seven pulses that lasted 30 seconds, 1 minute, or 3 minutes (Fig. 1B; modified from [30]). We exposed worms to one of 10 conditions: high or low concentrations of one of five chemical stimuli (Fig. 1C). Stimuli were either innately attractive or repellent as determined by previous chemotaxis and drop assays (Fig. 1C; [31, 32]). We monitored and tracked the activity of each neuron individually within a session, but we did not identify that same neuron across animals or sessions (i.e., Spontaneous, Buffer, Stimulus). Some neural responses were locked to the onset or offset of the stimulus in the Buffer and Stimulus sessions, and were likely sensory or interneurons involved in the detection and behavioral response to chemical stimulation [20]. Other neurons may be interneurons or motor neurons involved in motor commands [22]. A median of 44, 48, and 64 neurons were active during Spontaneous, Buffer, and Stimulus sessions, respectively, which indicated that more neurons were active during Stimulus sessions than either Spontaneous or Buffer sessions (Fig. 1D). All Stimulus sessions, however, activated similar numbers of neurons, regardless of stimulus identity (Fig. S1). Thus, network engagement increases in response to stimuli, and we next sought to assess whether different stimulus properties can be extracted using signatures of this change.

**Figure 1.**
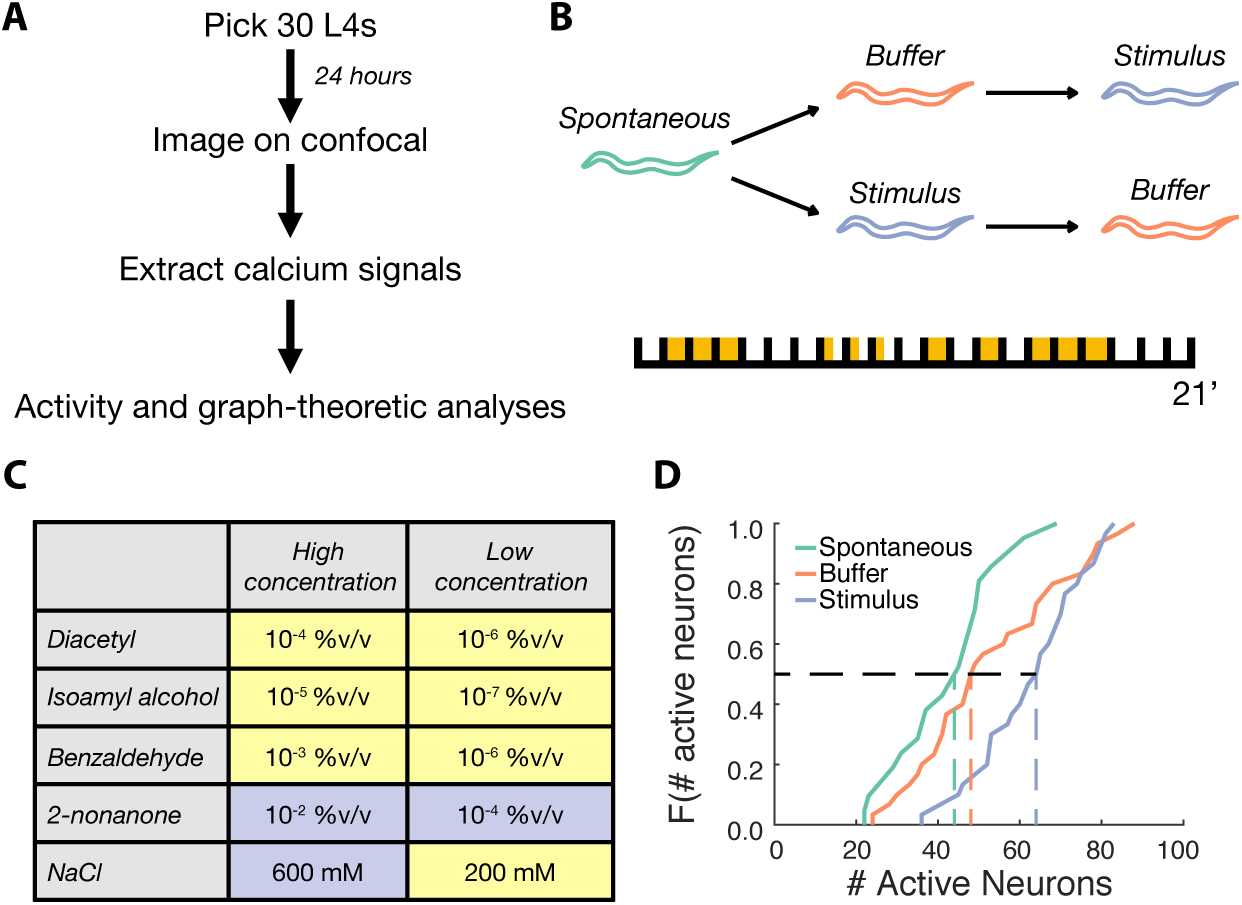
Whole-brain imaging experiments and analysis. A) Schematic showing the experimental protocol. Thirty stage L4 animals were picked onto a plate covered with OP50 24 hours before each experiment. Animals, as 1-day old young adults, were stimulated in an olfactory chip and their neural responses were imaged on a Zeiss Airyscan 880. Fluorescence traces were extracted from each video, and subsequently analyzed. B) Animals were imaged in three 21 minute-long sessions: Spontaneous, Buffer, and Stimulus. Buffer and Stimulus sessions used seven pulses of M9 buffer or one of 10 chemicals, respectively. C) Table showing the chemicals (attractants are colored yellow, repellents are violet) and concentrations tested. Three animals were tested per stimulus condition. %v/v refers to % vol/vol. D) The cumulative distribution function for the number of neurons active during Spontaneous (green), Buffer (orange), and Stimulus (blue) sessions shows that more neurons are active when a stimulus is present (p = 9.20E-6 and p = 4.61E-3 for the Spontaneous to Stimulus and Buffer to Stimulus comparisons, respectively, by the two-sample Kolmogorov-Smirnov test). N = 21, 30, and 30 animals for Spontaneous, Buffer, and Stimulus sessions, respectively.

### Stimulus valence and identity alter neural activity statistics

We first tested if simple measures of neural activity systematically changed with respect to different stimulus properties. We considered two ecologically-relevant stimulus properties: 1) Identity (diacetyl, 2-nonanone, benzaldehyde, isoamyl alcohol, or NaCl) and 2) Valence (attractive or repellent). We reasoned that an animal navigating its environment will recognize different chemicals (i.e., identity) as nutritious or toxic (i.e., valence). We focused on statistical measures (average and standard deviation of normalized neural activity) and measures that capture temporal dynamics (Fourier-based analysis of frequency spectra [33]). Each cell’s activity was normalized to the peak value it attained in the 21-minute session, and all measures were normalized to their pre-switch values.

We found that stimulus valence and identity have distinct effects on neural activity at stimulus onset (Figs. 2A,C, S2) but not offset (Figs. 2B,D, S3). For example, repellents (2-nonanone and 600mM NaCl) evoked higher mean neural activity than attractants (diacetyl, benzaldehyde, isoamyl alcohol, and 200mM NaCl) on stimulus onset (Figs. 2A, S2A), but not offset (Figs. 2B, S3A). The tastant NaCl induced more variable neural activity than the odorants on stimulus onset but not on offset (Figs. 2C,D, S2, S3), and there was no effect of stimulus identity on the mean activity (Figs. 2C,D, S2, S3). While repellents decreased the average frequency with the most power on stimulus onset (Figs. 2A, S2A), there was no difference between repellents and attractants on offset (Figs. 2B, S3A). For stimulus identity, the offset (Figs. 2D, S3B) of diacetyl and isoamyl alcohol slightly increased the peak frequency of neural oscillations (i.e., max of the frequencies with the most power in a sliding window over a 30-second period), but this effect was not observed on stimulus onset (Figs. 2C, S2B). Finally, there was no significant difference for most properties in buffer trials (Figs. 2, S2, S3), which indicates that these measures are sensitive to chemosensory stimulation. Thus, activity features are modulated by stimulus properties in an inconsistent manner at stimulus onset and offset.

Previous studies have shown that spontaneous activity in the nervous system of an immobilized worm lies in a low-dimensional PCA space [22, 23, 25]. Indeed, our analyses also suggest that, during the Spontaneous session, neural activity loops through a low-dimensional manifold, where the first three principal components (PCs) capture a median of 59% of the variance in our 21-minute long imaging sessions (Fig. 3A,C). However, we found that PCA was not able to infer different states during any type of stimulation – either Buffer or Stimulus changes – as both exhibited more complicated network dynamics than observed in an unstimulated worm (Fig. 3B). Specifically, the first three PCs captured a median of only 51% of the variance in stimulation sessions (Fig. 3C; p = 0.002 and 4.54E-5 for Spontaneous to Buffer and Spontaneous to Stimulus comparisons, respectively), and instead of smooth trajectories through PCA space, we observed jumps between different regions of state space (Fig. 3B). This agrees with [26], who found that a non-stimulated, but moving, worm also exhibits network dynamics that cannot be easily explained by the first three PCs.

**Figure 3.**
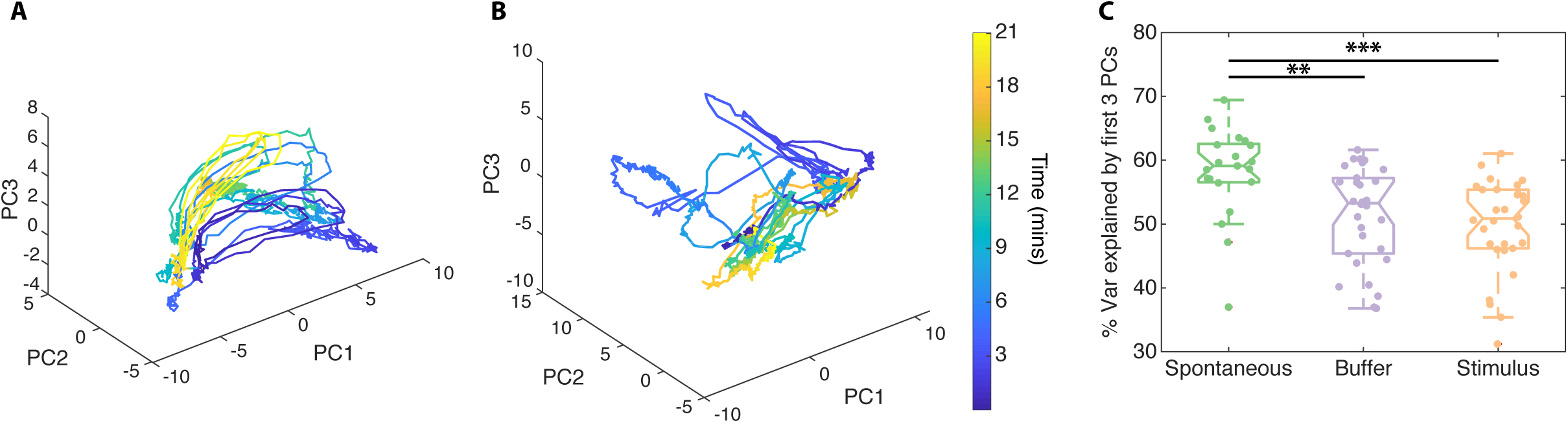
Observed neural dynamics preclude the use of PCA. Example neural dynamics observed during a Spontaneous session form loops through PCA-space (A), but not during a Stimulus session (B). Generally, the first three principal components explain a larger percentage of the variance during Spontaneous sessions than during Buffer or Stimulus sessions (C). Kruskall-Wallis test, with Dunn-Sidak post-hoc test, ** p < 0.01, *** p < 0.001.

Thus, a different approach is needed to decode stimulus properties from network dynamics in the absence of neural labels, particularly if traditional dimensionality-reduction-based methods fail to capture a significant portion of the variance. We next propose to use graph-theoretic features of the interactions between neurons.

### Computing graph-theoretic features of neural activity

We next asked if different stimulus properties could be better distinguished based on interaction patterns between neurons, instead of activity statistics of individual neurons. Graph-theoretic features capture complex interactions amongst groups of neurons. For example, these features can quantify how well a network can be divided into relatively independent clusters such that neurons in a cluster mostly interact with other neurons in the same cluster and sparsely interact with neurons in other clusters (this feature is called modularity). Other features capture how quickly information can spread through the network such that signals in one part of the network can be received and processed in another part of the network (one such feature is the largest eigenvalue of the graph) [34, 35]. Graph-theoretic features have been previously used to characterize activity changes in coarse brain networks (e.g., where nodes represent entire brain regions or large populations of neurons; reviewed by [34, 36]), but have not, to our knowledge, been used to analyze whole-brain activity at the single neuron level.

To characterize interactions between neurons, we first need to infer interactions between neurons based on their individual activities. For example, if neuron B’s activity rises shortly after neuron A’s activity rises, then we may infer a functional interaction between neurons A and B. Formally, to predict interactions, we computed the normalized mutual information (NMI; [37]) between the activity vectors of every pair of neurons in a 30-second period around a stimulus switch (i.e., onset or offset). NMI measures how much information one variable contains about another variable, which in this context reveals a putative interaction between the activity vectors of two neurons (see Materials and Methods). We computed the NMI between all pairs of *n* neurons in a given worm during a 30-second period of interest (either before or after a stimulus switch). We focused on a 30-second period for two reasons: 1) an animal can begin to move toward or away from an attractant or repellent, respectively, well within 30 seconds [30, 38], and 2) sensory neurons that detect chemical stimuli tend to reach their maximum response within ∼10 seconds of stimulus onset [20]. Thus, analyzing 30 seconds following a stimulus switch should be sufficient to capture the representation of ecologically-relevant information in neural activity [20, 32, 38].

We then computed a weighted graph *G=(V,E)*, where *V* is the set of nodes (neurons) and *E* is the set of weighted edges (inferred functional interactions) between neurons. Each edge weight equals the NMI between the two neurons, which lies between 0 and 1, where a larger number implies a stronger interaction. Pearson’s correlation (PC) has been previously used [39] to generate functional connectivity networks with both positive and negative weights (as well as 0 for uncorrelated neurons, which is rare in practice); the former indicates that both neurons increase or decrease their activity together (e.g., excitation), and the latter implies that as one neuron increases its activity, the other neuron’s activity decreases (e.g., inhibition). As a group, GCaMPs are known to faithfully reflect increases in a cell’s internal calcium concentration, and thus its excitation; however, GCaMPs were not optimized to reflect a decrease in calcium concentration, or its hyperpolarization (e.g., see [40, 41]). Further, many graph-theoretic analyses require that all edge weights be non-negative, and we found that 46.8% of edges had a negative weight when we used Pearson’s correlation. Thus, we did not use weights from Pearson’s correlation, though we did use it in separate analyses to independently study the putatively excitatory and inhibitory subnetworks.

From each graph, we extracted network features that capture different interaction patterns at both the local and global network scales. Specifically, we focused on five classes of network measures, including basic structure, functional segregation, functional integration, centrality, and resilience (summarized in Table S1 and [34]). Basic structure refers to general aspects of the graph, such as the largest eigenvalue or the median weight of the network, which indicates how strongly nodes interact with one another. Functional segregation measures how much processing occurs in small groups of neurons, or modules, and encompasses the network’s modularity and number of modules – in a more modular network, neurons cluster into groups that strongly communicate with one another. Functional integration indicates how efficiently different groups of neurons can pass information to each other; a representative measure is the average shortest path distance between pairs of neurons in the network, which indicates how quickly, on average, information can pass between any two neurons. Centrality measures how important, or central, a neuron is to information transmission between different parts of the network and can be assessed, for example, using the average betweenness centrality – the average fraction of shortest paths linking any two neurons that pass through a given neuron. Finally, measures of resilience to perturbations, such as lesions, includes the average assortativity coefficient – the average correlation coefficient between the degrees of any two connected neurons.

For fair comparisons, each graph-theoretic feature was normalized to account for any dependence on network size [42]. Further, to highlight changes in a graph-theoretic property after a stimulus switch, we used the same normalization scheme we used for neural activity features – in short, we divided the post-switch value of the feature to its pre-switch value. This was a critical normalization scheme as the organization of each worm’s neural network was quite variable, even in the absence of stimulation (Fig. S4). Thus, we report how the addition or removal of a stimulus affected patterns of neural network activity.

### Graph-theoretic features distinguish stimulus identity

For each stimulus property, we tested the extent to which any graph-theoretic feature significantly changed in response to a stimulus switch. Overall, some network features showed a reliable change upon both switches (onset and offset), whereas activity features of neurons revealed no such reliable change. For example, the average betweenness centrality reliably changed with respect to stimulus valence. Repellents decreased the network’s average betweenness centrality on both stimulus onset and offset, while attractants had no effect (Figs. 4A,B; Table S2). Furthermore, these changes were not concomitant with a change in the median weight of the network (Fig. 4A,B; Table S2), which indicates that the change in average betweenness centrality is not simply driven by stronger or weaker interactions in one condition over another, but rather by differences in the patterns of interactions amongst neurons. Thus, valence modulates the centrality of neurons bridging disparate regions of the network. Surprisingly, these effects occur both at stimulus onset and offset, indicative of a fundamental signature in how each class of stimulus is processed in the *C. elegans* nervous system.

**Figure 4.**
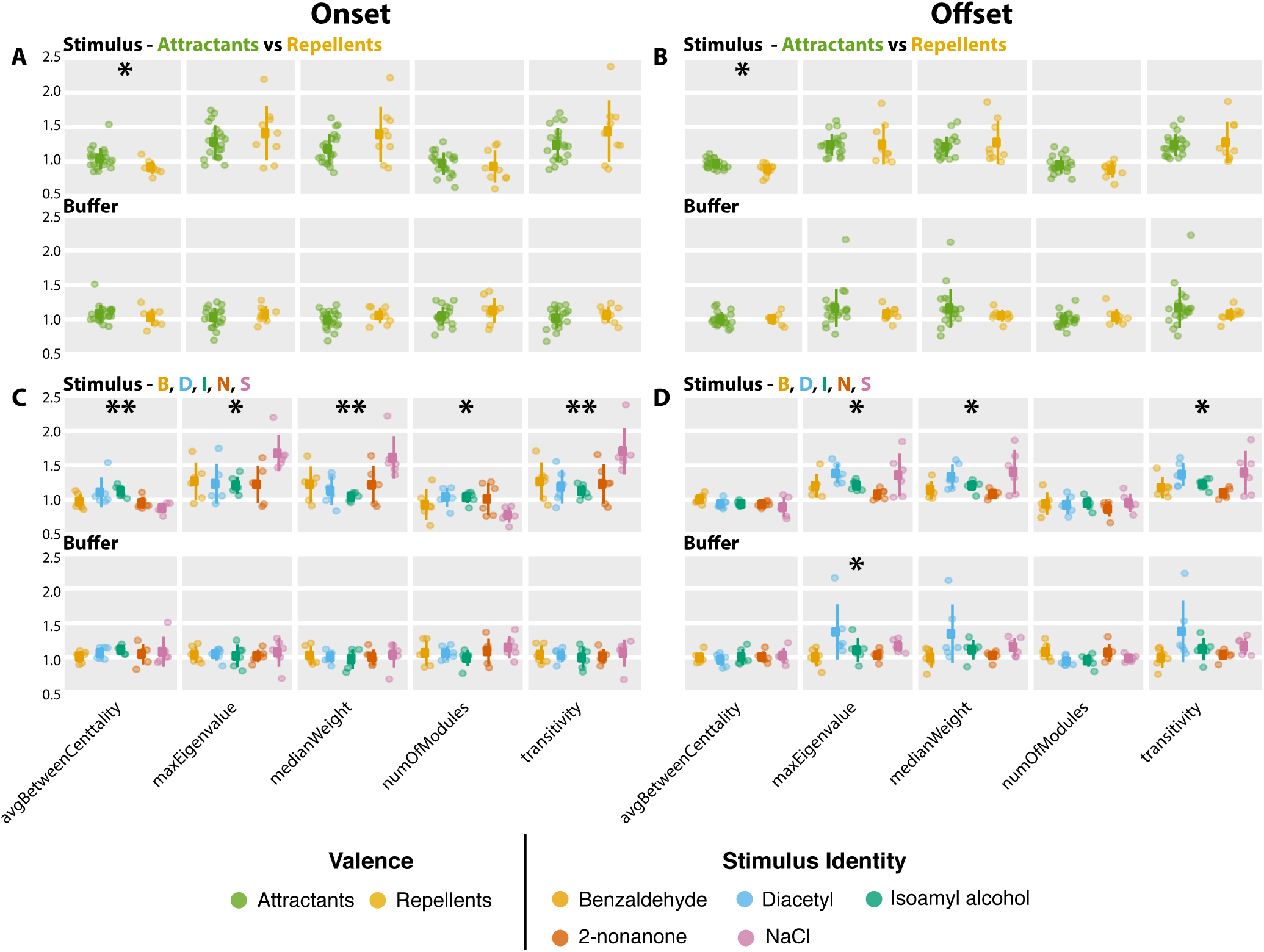
Stimulus valence and identity have distinct effects on network features. Repellents consistently induce a decrease in average betweenness centrality on stimulus onset (A) and offset (B). Stimulus identity also affects the average betweenness centrality on stimulus onset (C), but not offset (D), in addition to effects on various other network features. There were no changes in any of the measured properties on buffer onset (A, C) or offset (B, D). N = 21 for attractants and N = 9 for repellents (A, B), and N = 6 for each chemical stimulus (C, D). * p < 0.025, ** p < 0.005, by likelihood ratio test on full and null generalized linear mixed-effects model, where the former included either valence or identity as a fixed effect.

Stimulus identity affected several features, though there were only a few features - the median network weight, transitivity, and max eigenvalue - that reliably changed on both stimulus onset and offset (Figs. 4C,D, S5, S6; Table S2). When worms were exposed to NaCl, the network had the largest median weight and the smallest diameter (i.e., the furthest distance between any two neurons). This indicates that the NaCl-induced network could efficiently transmit information across the network. Furthermore, this network had the largest transitivity (i.e., average strength of interactions between connected triplets of neurons) and max eigenvalue (the larger the max eigenvalue, the more easily signal spreads through a network), suggesting a unique combination of strong local connectivity and efficient global reach, which is often observed in small-world networks [43]. Buffer sessions modulated no graph-theoretic features on buffer onset (Figs. 4, S5, S6; Table S2); however, for stimulus identity, buffer offset did effect change in several graph-theoretic features, but not median weight (Fig. S6; Table S2). This indicates that changes in median weight may require a stimulus, consistent with previous studies showing that stimuli drive correlated activity across multiple neurons [44–46]. Overall, stimulus properties affect network connectivity in a reliable manner, and these features may serve as decoding signatures.

Graph-theoretic features quantitatively capture differences in network structures that emerge during stimulus presentation. To illustrate these differences more intuitively, we visualized how the networks changed shape before and after stimulus onset. Figure 5 shows the network of one worm before the onset of 200 mM NaCl (Fig. 5A,C), and during the first 30 seconds of the pulse (Fig. 5B, D). In the latter, there is an increase in the number of strong edges (i.e., edges of weight larger than 0.2), a decrease in number of modules (from 7 to 5), and a decrease in the betweenness centrality of several neurons. More neurons are connected with the rest of the network (thereby increasing the max eigenvalue), and triplets of neurons are more likely to be strongly connected (i.e., a larger transitivity). Due to the normalization scheme employed, changes in the values of graph-theoretic features may appear to be mild; however, as illustrated, these changes capture significant differences in connectivity patterns between neurons and may be biologically informative.

**Figure 5.**
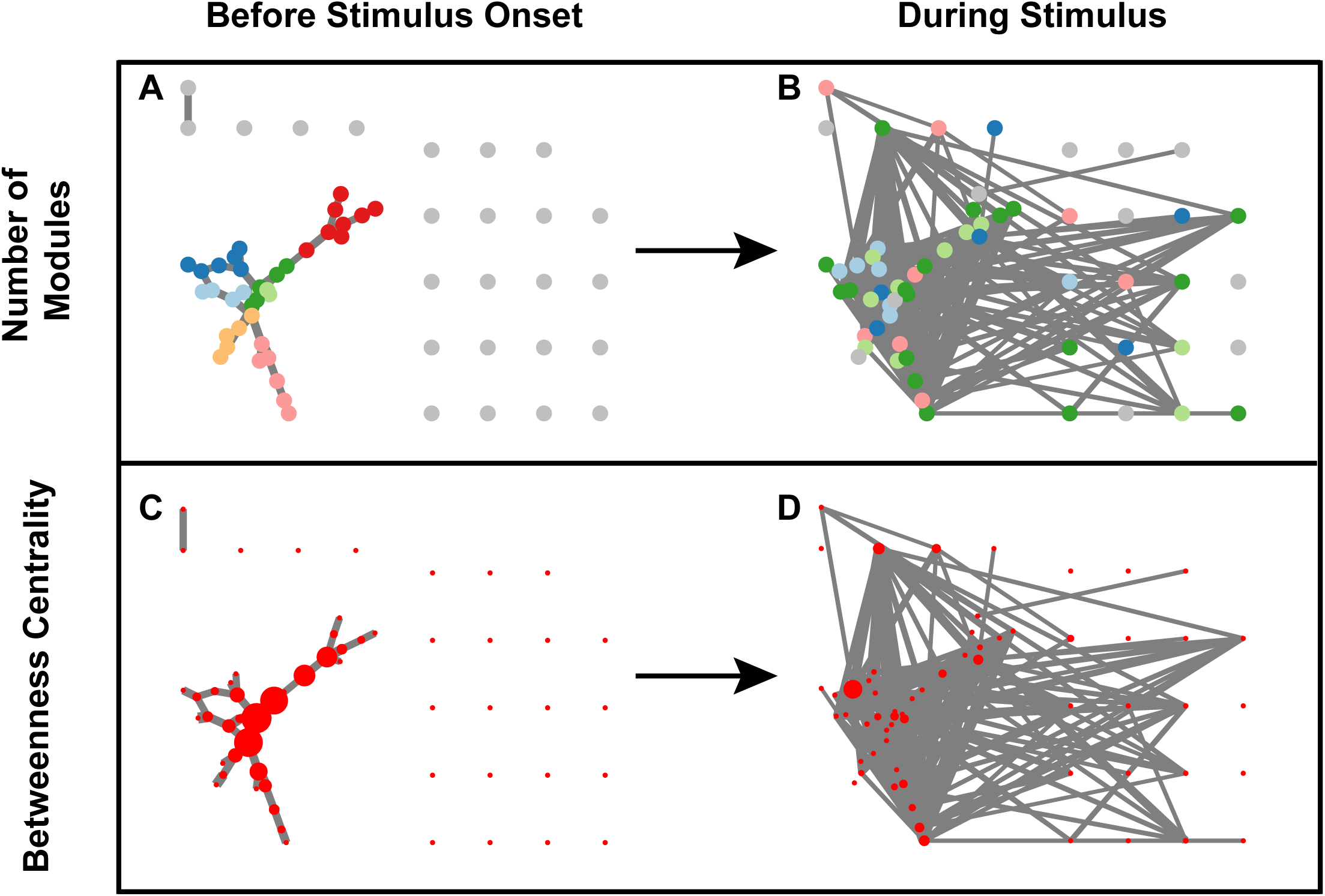
NaCl induces a decrease in the number of modules and average betweenness centrality of the worm’s neural network. The networks depicted here, for one example worm, show neurons (circles) connected by lines (edges); the thickness of the edges correspond to the amount of normalized mutual information between the two linked neurons. Each neuron’s position is fixed across all four panels. Prior to stimulation with 200 mM NaCl (A, C), this worm’s functional connectivity had 7 modules (A) and fairly central neurons (C). During the first 30 seconds of stimulation, however, the number of modules dropped to 5 (B has fewer colors than A), with fewer central neurons (D has smaller circles than C). Note the increase in recruited neurons during (B, D), relative to before (A, C), stimulus. Neurons with the same color in A and B belong to the same module. The size of the neurons in C and D are positively correlated with their betweenness centrality. All edges less than 0.2 were removed for the purpose of visualization. Thus, isolated neurons only had weak edges to all other neurons.

### Using machine learning methods to predict stimulus properties

We used a machine learning approach to test how well neural activity features (Fig. 2) and graph-theoretic features (Fig. 4) could predict stimulus properties on the first stimulus pulse, when the animal has not undergone any adaptation. As before, the stimulus properties we considered were stimulus valence and identity. We used a logistic regression classifier, which is a simple and commonly used classifier that has no built-in assumptions about the distribution of the data. We evaluated its ability to generalize to unseen networks using cross-validation, followed by a permutation test to assess its performance against an empirically-derived chance level of accuracy (see Materials and Methods). We also combined graph-theoretic and neural activity features, to see if graph theory and neural activity provided distinct information that could collectively improve classification accuracy.

**Figure 2.**
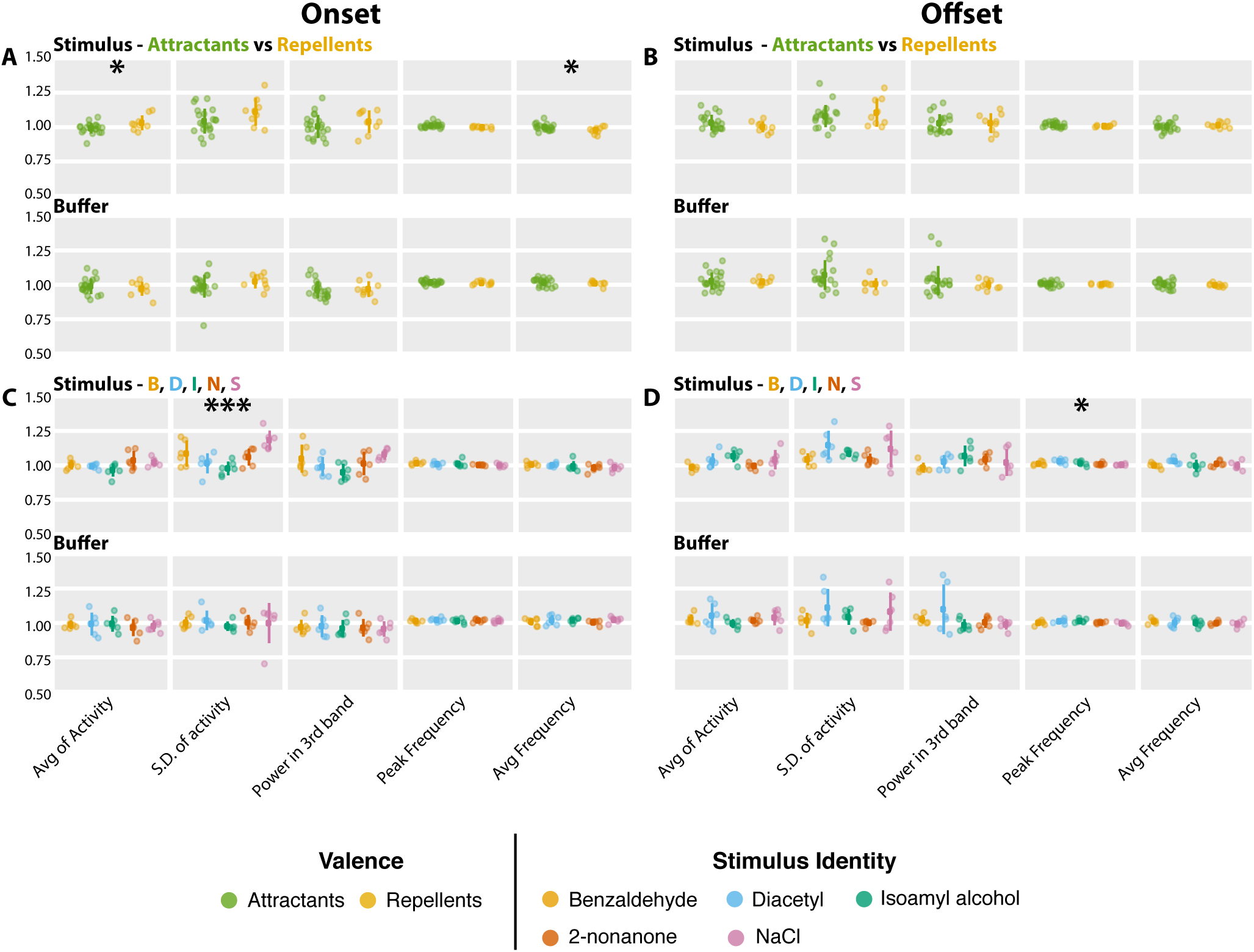
Stimulus valence and identity have distinct effects on neural activity. Repellents induce an increase in average neural activity and a decrease in the average frequency of the peak frequency on stimulus onset (A), with no changes on stimulus offset (B). On the other hand, stimulus identity affects the variability (i.e., standard deviation) of neural activity on stimulus onset (C), and the peak frequency on stimulus offset (D). There were no changes in any of the measured properties on buffer onset (A, C) or offset (B, D). Avg and S.D. of activity refer to average and standard deviation of neural activity. Power in 3rd band refers to average power in the frequency range from 0.34 - 0.47 Hz. Peak frequency is the frequency with the most power in a 30-second bin, and avg frequency is the average of the frequencies with the most power in a sliding-window bin covering a 30-second period. N = 21 for attractants and N = 9 for repellents (A, B). N = 6 for each chemical stimulus (C, D). * p < 0.025, *** p < 0.0005, by likelihood ratio test on full and null generalized linear mixed-effects model, where the former included either valence or identity as a fixed effect.

The logistic regression classifier performed well on predicting stimulus identity on onset, but not offset (Fig. 6B,D, and Table S7). In particular, accuracy was high when using graph-theoretic features alone (40% accuracy, chance: 20% permutation accuracy: 16±8%, p-value = 0.004). Adding neural activity features increased the accuracy slightly (47% accuracy, Fig. 6B). On the other hand, when training the classifier using only neural activity data, accuracy was far lower (17%; Fig 6B). Finally, we did not exceed chance accuracy when classifying attractants and repellents using either of the three feature sets (Fig. 6A,C, and Table S7), or any of the buffer sessions except for identity on buffer offset (Fig. S7D and Table S7). However, buffer sessions that preceded stimulus sessions did not show above chance accuracy (Fig. S8), suggesting that the above-chance performance on all buffer sessions combined might be confounded by prior stimulus experience. Thus, graph-theoretic features alone appear capable of quantitatively discriminating identity on stimulus onset, with some gain when also including measures of neural activity in combination. Together, these data show that the *C. elegans* neural network responding to stimulus, but not buffer, has structural characteristics that can be identified using graph theory with a simple machine learning classifier.

**Figure 6.**
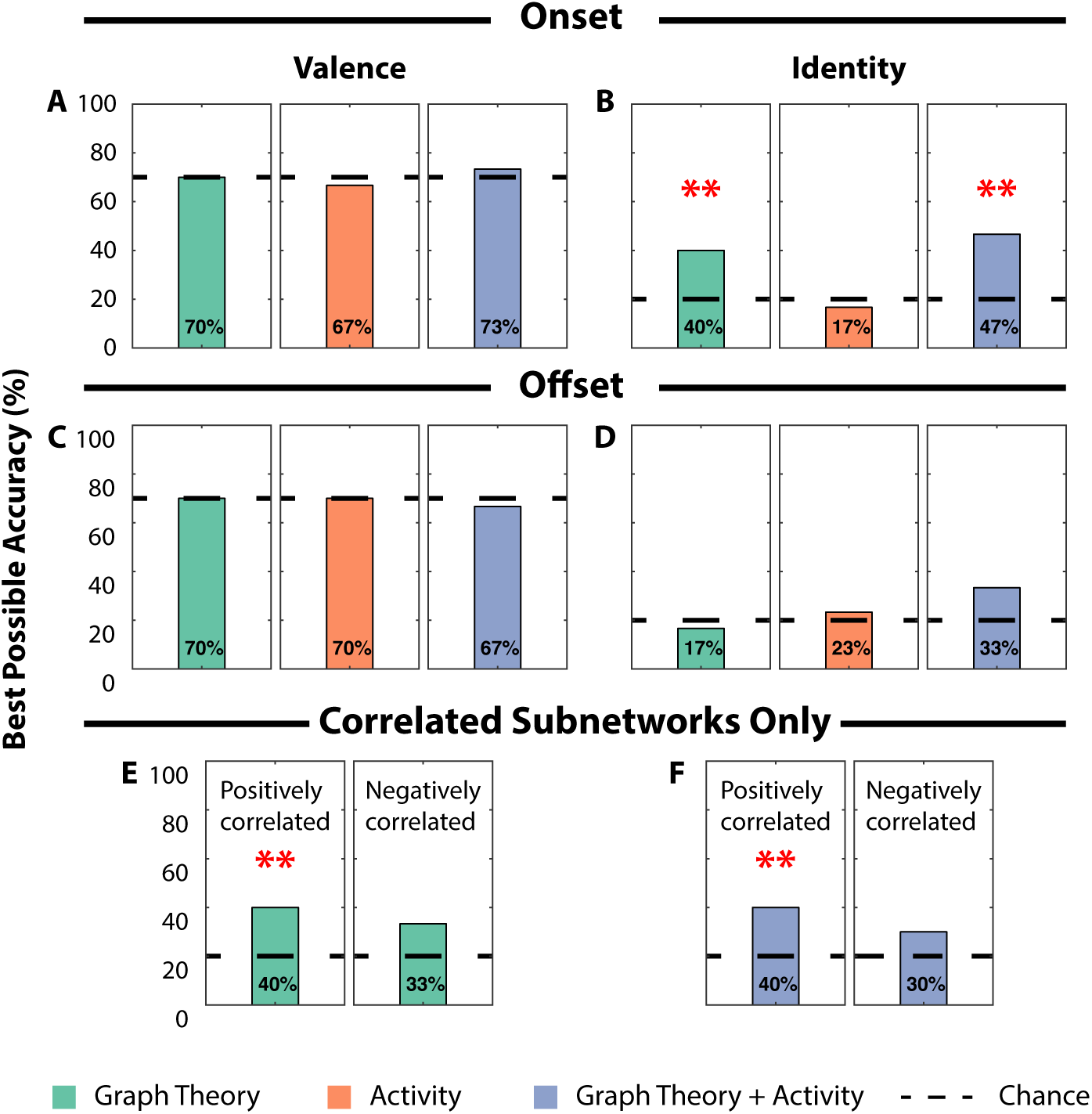
Network features increase classification accuracy of stimulus identity. Classification accuracy achieved on the first pulse of Stimulus sessions using a Logistic Regression classifier trained using network features (green), neural activity features (orange) or both (blue). Accuracies are shown for stimulus valence and identity at stimulus onset (A, B) and offset (C, D), respectively. The accuracy on correlated subnetworks for stimulus identity at stimulus onset is shown for only network (E) or combined features (F). Red asterisks indicate significantly above-chance classification accuracy as determined with a permutation test (**p < 0.005). Black dotted lines indicate theoretical chance accuracy.

### Contributions of putative excitatory and inhibitory subnetworks to discrimination of stimulus identity

NMI can detect non-linear interactions between two signals but is unable to determine the sign of the interaction. For instance, two neurons can have high NMI, yet they may be positively or negatively correlated. The sign of the correlation is important because it implies different types of interactions, both of which are biologically meaningful – positive correlations are thought to indicate excitation, while negative correlations may imply inhibition, although this is not always true (e.g., if neurons A and B each receive input from a common, third neuron, then A and B will be positively correlated, in spite of the lack of a direct interaction between them). To identify the contributions of the positively and negatively correlated subnetworks to discriminating stimulus identity, we computed adjacency matrices based on Pearson’s correlation and used the sign of those networks to split the NMI-based networks into two subnetworks: one composed of putatively excitatory interactions, and another composed of putatively inhibitory interactions. This enabled us to capture all non-linearities in neuronal interactions and analyze each subnetwork independently.

Unlike our results on the full network, we found no consistent differences between attractants and repellents on stimulus onset or offset for either the positively or negatively correlated subnetworks alone (Table S6). This suggests that the decrease in average betweenness centrality seen in the full network (Fig. 4A,B) is driven by a decrease in the centrality of neurons that integrate both excitatory and inhibitory signals. On the other hand, several network features consistently changed on stimulus onset and offset for both subnetworks in an identity-dependent manner (Table S6). While not all of these features are shared between the excitatory and inhibitory subnetworks, many, such as the network diameter and median weight, are. Finally, similar to the full network, there are few valence or identity-dependent changes on buffer onset, and a few on buffer offset (Table S5). Curiously, many of these identity-dependent changes are only seen in the negatively-correlated subnetworks, which indicates that the inhibitory subnetwork may carry a stronger memory of past stimulus.

Finally, we tried to classify stimulus valence and identity using network features solely derived from one subnetwork at a time. We found that the network features based on the positively correlated subnetwork contained as much discriminatory power as network features based on the full network (40% accuracy for both, chance: 20% permutation accuracy: 14±8%, p-value = 0.003; Fig. 6E,F, and Table S11), while the negatively correlated subnetwork’s features contained no discriminatory power (Fig. 6E,F, and Table S12). This implies that stimulus identity most strongly modulates the pattern of excitation, but not inhibition. Oddly, neural activity features did not change the discriminatory power for the positively correlated subnetwork features (Table S11), which suggests that the interplay of excitation and inhibition together allows the context of neural activity to improve accuracy.

Overall, these additional analyses further demonstrate the value of using graph-theoretic features to decode stimulus properties. We anticipate that other parameters introduced when extracting features (e.g., modifying the size of time windows, inferring directionality, measuring the perceived intensity of a stimulus) may provide further biological insight.

## Discussion

A central challenge in neuroscience is to develop tools that can uncover how stimulus properties, such as valence and identity, are encoded within neural networks. We monitored changes in the activity of over 40 neurons in the head of *C. elegans* as it experienced diverse chemical stimuli. We found that some activity statistics, such as the mean and standard deviation of neural activity and Fourier-based frequency measures, are modulated by, yet unable to, distinguish stimulus valence and identity. We then extracted graph-theoretic features from time-series activity traces of neurons and found improved ability to predict stimulus identity. Our results suggest that some stimulus properties can be decoded using network features, which could be useful in instances where neuron labels are unknown or difficult to map across animals.

We used normalized mutual information to generate functional connectivity networks from neural activity traces. In contrast, recent work has tried to infer causal networks from activity traces using methods such as adaptive, sparsity-constrained Granger causality [47], network deconvolution [48], convex risk minimization [49], and convergent cross mapping [50]. Some of these methods, however, make assumptions about the data type and nature of causality that may not be true here; e.g., they assume spiking data as opposed to graded potentials, or they assume specific models of information spread, such as those used in epidemiology. While we used a relatively simple approach, our method is general and allows for any network inference algorithm to be plugged into our framework.

The median network weight, transitivity, and max eigenvalue all emerged as useful features for discriminating stimulus identity. This implies that neural networks likely use changes in strength of interactions, especially amongst local triplets of neurons, to facilitate widespread communication and process stimulus identity. Consistently, amongst the five stimuli tested, NaCl always attained the greatest increase in median weight, max eigenvalue, and transitivity, which may reflect its ability to use both salt and odor circuitry to encode sensory information [17]. The rest of the chemicals induced similar increases in the max eigenvalue and transitivity of the nematode neural network on stimulus onset, yet had more differentiated responses on stimulus offset, which indicates that the initial detection of an odorant might evoke core changes in neural network organization, supplemented with additional chemical-specific changes.

Stimulus valence, on the other hand, modulated the network’s average betweenness centrality. While repellents led to network activity with a smaller average betweenness centrality, attractants did not produce substantial changes in centrality. This implies that repellent information is likely processed by fewer central neurons bridging disparate parts of the network, while attractants may require a large number of neurons across the entire network. Consistently, previous studies have shown that attractants are typically encoded by a combination of multiple sensory neurons [18], while the onset and offset of the repellent 2-nonanone is detected by the AWB chemosensory neurons [51].

While we chose several popular graph-theoretic features to study diverse aspects of network organization [34, 52], this list is not exhaustive, and there may be other features that can better discriminate stimulus properties. We also focused on fully-connected weighted networks, as opposed to sparse networks by applying an arbitrary threshold to remove weak edges. Sparse network analysis depends on the precise threshold and might result in data that is difficult to interpret [53]. Consistently, attempts to use three different proportional thresholds to filter network weights showed that different features emerged as significant depending on the threshold chosen (Tables S3, S4), and the ability to classify stimulus identity was similarly variable (Tables S8 – S10). Finally, we used a basic machine learning approach to quantitatively test the power of using network features to classify stimulus properties. While we both used sample sizes (n=30 animals) comparable with prior studies and adopted standard approaches to avoid over-fitting (e.g., cross-validation, permutation testing), we nonetheless caution that building reliable machine learning models with relatively small datasets can be sensitive to a few data points. As it becomes experimentally easier to generate larger datasets with more neurons across many more conditions in other species, we hope our framework of using graph-theoretic features of network activity to understand neural function can be further improved.

## Materials and Methods

### Whole-brain calcium imaging

All imaging experiments were performed on a previously published strain (ZIM1048 *lite-1(ce314)X, msmIs4* [23]). We trapped 30 young adults in a modified olfactory chip [19] that orients animals similarly [21]. Changes in GCaMP fluorescence were monitored using a Zeiss LSM 880 Airyscan (1.27-1.62 volumes/second) while the animal’s nose experienced buffer or one of five stimuli (Diacetyl 10^-4^ %vol/vol, 10^-6^ %vol/vol [54, 55], benzaldehyde 10^-3^ %vol/vol, 10^-6^ %vol/vol [18], isoamyl alcohol 10^-5^ %vol/vol, 10^-7^ %vol/vol [20, 56], 2-nonanone 10^-2^ %vol/vol, 10^-4^ %vol/vol [55] and NaCl at an aversive 600mM [32] and at an attractive 200mM [57] concentration). For each animal, we obtained a 21-minute recording with no stimulation (“Spontaneous”), a second 21-minute recording with M9 buffer changes (“Buffer”) and a third 21-minute recording with stimuli changes (“Stimulus”). Buffer and Stimulus sessions were interleaved for different animals except for those experiencing 2-nonanone, when Buffer always preceded Stimulus. The Stimulus or Buffer pattern was adapted from [30].

### Data processing

We first deconvolved the pixels using built-in Airyscan processing tools. We then corrected for motion using NormCorre (https://github.com/flatironinstitute/NoRMCorre) and extracted the raw fluorescence traces using CaImAn (https://github.com/flatironinstitute/CaImAn-MATLAB) [58–63]. We used multiple iterations using grids of decreasing size to analyze high concentration NaCl “Stimulus” trials, which had larger motion artefacts compared to other stimuli. Importantly, the low concentration NaCl “Stimulus” trials did not require this extra preprocessing step. Nine of thirty “Spontaneous” sessions had too much movement to correct, and were not further analyzed. To accurately identify all neurons in the head of the animal, we set the cutoff K, for the number of components to look for, to 140 (this is higher than the number of neurons previously detected in this strain [23]) allowing the CaImAn algorithm to detect many neurons with a high signal-to-noise ratio as recommended by [63]. Hence, neurons with no change in fluorescence were not detected. Finally, we used custom software to manually verify that regions of interest extracted by our analysis pipeline qualitatively matched the video, and to eliminate non-neuronal ROIs (e.g., gut granules).

### Measuring statistical features of neural activity

Each neuron’s activity trace was normalized within a session by the max fluorescence value reached in that 21-minute imaging session. To highlight the change in neural activity properties after a stimulus switch, we normalized the value of the property to its value pre-switch. For example, for stimulus onset, we computed the feature in the first 30-seconds after stimulus onset and divided by the value of the feature in the 30-seconds prior to stimulus onset. Thus, we report how much stimulus onset or offset changed the value of the feature from its pre-switch baseline to reflect how the addition or removal of a stimulus affected neural activity.

All analyses of the temporal dynamics of neural activity was performed using Fourier transforms. We used MATLAB’s periodogram to average the power of the four frequency bands used in our analyses (1^st^ band: 0.07 – 0.2 Hz, 2^nd^ band: 0.2 – 0.33 Hz, 3^rd^ band: 0.33 – 0.47 Hz, 4^th^ band: 0.47 – 0.6 Hz). We used MATLAB’s spectrogram to determine the max, average, and standard deviation of the frequency with the most power in a 30-second bin, using 10-second long sliding windows with 50% overlap. Thus, our measures quantified how much a switch from buffer to stimulus or stimulus to buffer changed activity.

### Determining functional connectivity

For each session, every neuron’s activity was normalized by the peak value it reached in the entire 21-minute imaging session, bounding every neuron’s activity in the range [0, 1]. Each session generated an *n* x *T* matrix, where *n* is the number of neurons and *T* is the length in time of imaging. Using this matrix, we generated two adjacency matrices, one for the pre-switch period and one for the post-switch period. Each of these periods lasted 30 seconds and resulted in 28 periods of activity (each session had 7 pulses, 2 switches per pulse, and 2 pre- and post-switch periods per switch). We made two types of adjacency matrices for every block: one based on the absolute value of the Pearson’s correlation (PC), and the other based on the normalized mutual information (NMI) with a 0.1 bin size. The former was calculated using MATLAB’s built-in corr function. The latter was calculated per [37] as: 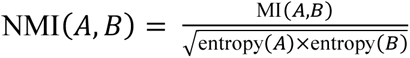, where entropy(A) is the entropy of the activity of neuron A, and MI(A,B) is the mutual information between the activity patterns of neurons A and B. While many variants of NMI exist [64], the one we used has the desirable property of only reaching its maximum value when the distributions of states of two neurons are identical. If a neuron had an entropy of 0 because all its activity during the entire 30-second period was localized to one histogram bin (e.g., 0-0.1), then its edges to all other neurons were set to 0, effectively decoupling the neuron from the network; this only occurred for 7.3%, or 8,930 out of 123,116 cells across all imaging sessions and worms. We reasoned that a neuron whose activity did not appreciably change in a 30-second time period is not likely to be involved in processing of that stimulus. Finally, all self-loops were removed by setting the diagonal of the adjacency matrix to 0.

### Graph-theoretic analysis

Graph-theoretic features were calculated using the Brain Connectivity Toolbox [34, 52] on the largest connected component of each network. For features that produced distributions, such as local efficiency, we report the mean value. The degree distribution, radius, diameter, largest eigenvalue, and average shortest path, were all normalized by the number of neurons in the largest connected component. We used the Louvain community detection algorithm to find modules [65].

### Machine learning classification

The logistic regression classifier was built using one of three sets of features. The first set was composed of graph-theoretic features. The second set (called activity features) included the mean and standard deviation of neural activity. The third set of features combined the first and second sets. All graph-theoretic features used to classify stimulus valence and identity came from adjacency matrices constructed using NMI. A logistic regression classifier was trained on these three sets of features, and built using Python’s scikit-learn library [66], with default settings and using leave-one-out cross-validation to test generalization error. Finally, to avoid over-fitting, we used permutation testing (Test 1 in [67]) in which we randomly permutated the labels of the classes 1000 times and compared each classifier’s performance to its own null distribution. We took the relative position of the classifier’s true accuracy (trained on non-permutated labels) in this permutation distribution to be its p-value [67, 68]. This was a critical test that ensured our results were not merely due to chance, which is a significant yet often unappreciated issue in machine learning applications because theoretical chance is defined for sample sizes that are infinitely large [68].

### Statistical analyses for mixed-effects models

Our experiments consisted of multiple measurements from the same animal, grouped according to three ecologically-relevant aspects of chemical stimuli. This repeated-measures design is often modeled using a mixed-effects model because it accounts for: 1) fixed-effects owed to the ecologically-relevant condition (e.g., valence), and 2) random-effects due to animal variability (e.g., some animals might naturally have a larger modularity). Thus, we can provide a mixed-effects model with a matrix that contains information on the time in seconds since the first pulse, graph-theoretic features assessed at that pulse, and an indicator variable denoting which stimulus property the animal experienced (e.g., attractant or repellent). A mixed-effects model that better explains the data using the class indicator than without it, tested using a likelihood-ratio test, indicates a significant contribution from the fixed-effect to the model fit. We used linear mixed-effects models when the data were qualitatively normally distributed, as tested with a quantile-quantile plot using MATLAB’s qqplot, or a generalized linear-mixed effects model with a response distribution modeled as a gamma distribution otherwise.

### Availability of data

Data and code to reproduce all analysis steps and figures will be made available at: https://osf.io/z4aq3/.

## Acknowledgements

We thank U. Manor, T. Zhang and Z. Cecere for technical help with our imaging experiments; C. Lee-Kubli and M. Rieger for help with statistics; and M. Matty, C. Lee-Kubli, J. Haley, M. Rieger, K. Quach, A. Chandrasekhar and J. Fleischer for helpful comments on the manuscript. This work was supported by grants from the Pew Charitable Trusts, NIH 1R01DC017695 (S.N.) and NIH 5R01MH096881 (S.H.C). J.J.H was supported by a Graduate Research Fellowship (DGE-1650112) from the National Science Foundation, Jesse and Caryl Philips Foundation and by NIH F99NS115336.

## Supporting Information

**Supplemental Figure S1.**
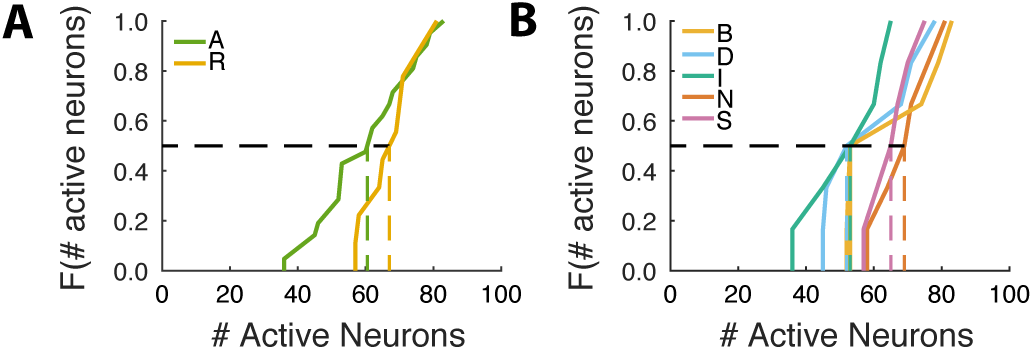
Chemical classes do not activate different numbers of neurons. The cumulative distribution of number of active neurons does not differ in a (A) valence- or (B) identity-dependent manner. All p > 0.05, by two-sample Kolmogorov-Smirnov test. N = 21 for attractants and N = 9 for repellents (A), and N = 6 for each chemical stimulus (B).

**Supplemental Figure S2.**
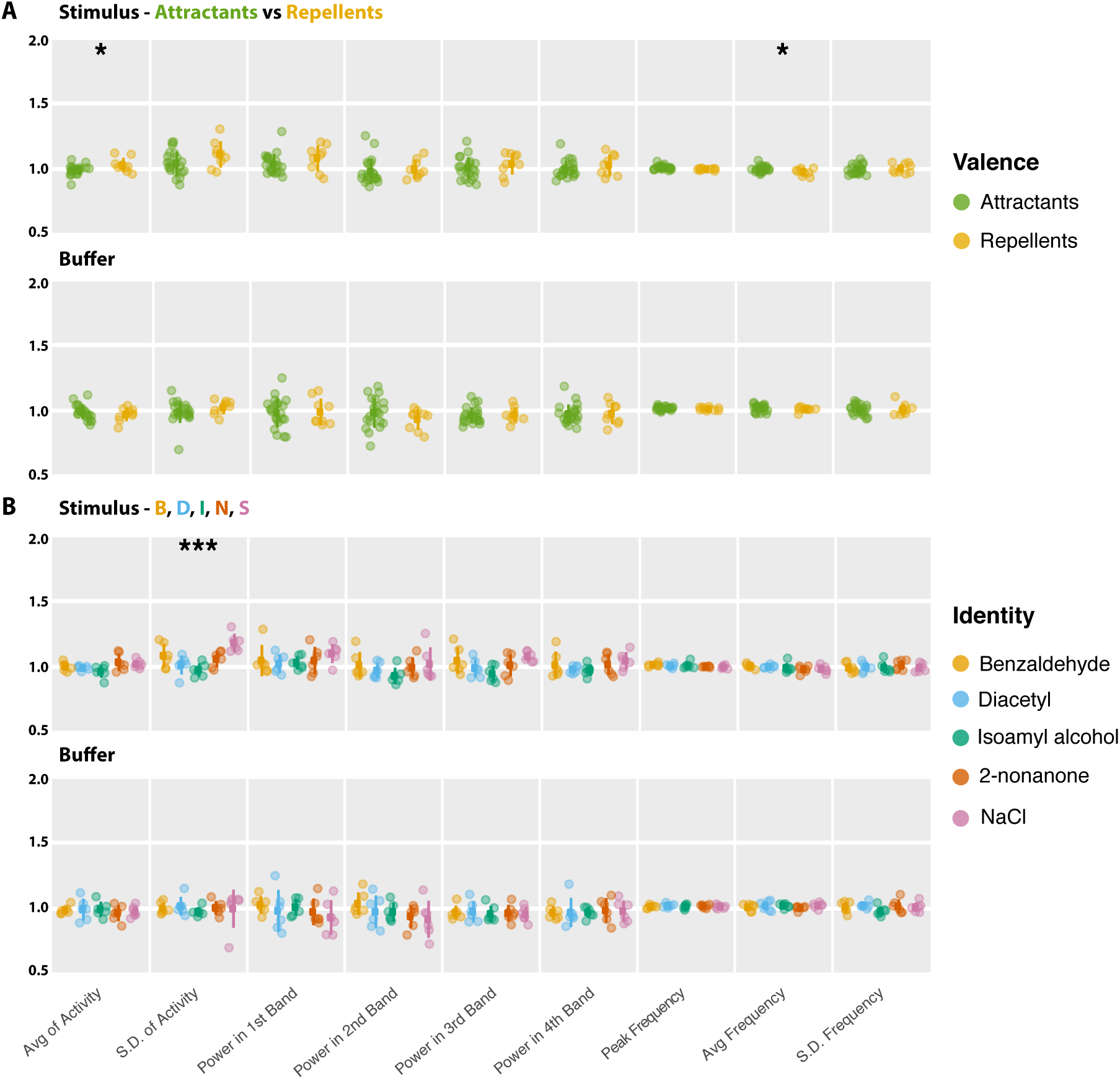
Stimulus valence and identity have distinct effects on neural activity on stimulus onset. Repellents induce an increase in average neural activity and a decrease in the average frequency of the peak frequency on stimulus onset (A). On the other hand, stimulus identity affects the variability (i.e., standard deviation) of neural activity on stimulus onset (B). There were no changes in any of the measured properties on buffer onset (A, B). Avg and S.D. of activity refer to average and standard deviation of neural activity. Power in 1st, 2nd, 3rd, and 4th bands refer to average power in the frequency ranges from 0.07 - 0.2 Hz, 0.2 - 0.34 Hz, 0.34 - 0.47 Hz, and 0.47 - 0.6 Hz. Peak frequency is the frequency with the most power in a 30-second bin, and avg frequency and S.D. frequency are the average and standard deviation, respectively, of the frequencies with the most power in a sliding-window bin covering a 30-second period. N = 21 for attractants and N = 9 for repellents (A), and N = 6 for each chemical stimulus (B). * p < 0.025, *** p < 0.0005, by likelihood ratio test on full and null generalized linear mixed-effects model, where the former included either valence or identity as a fixed effect.

**Supplemental Figure S3.**
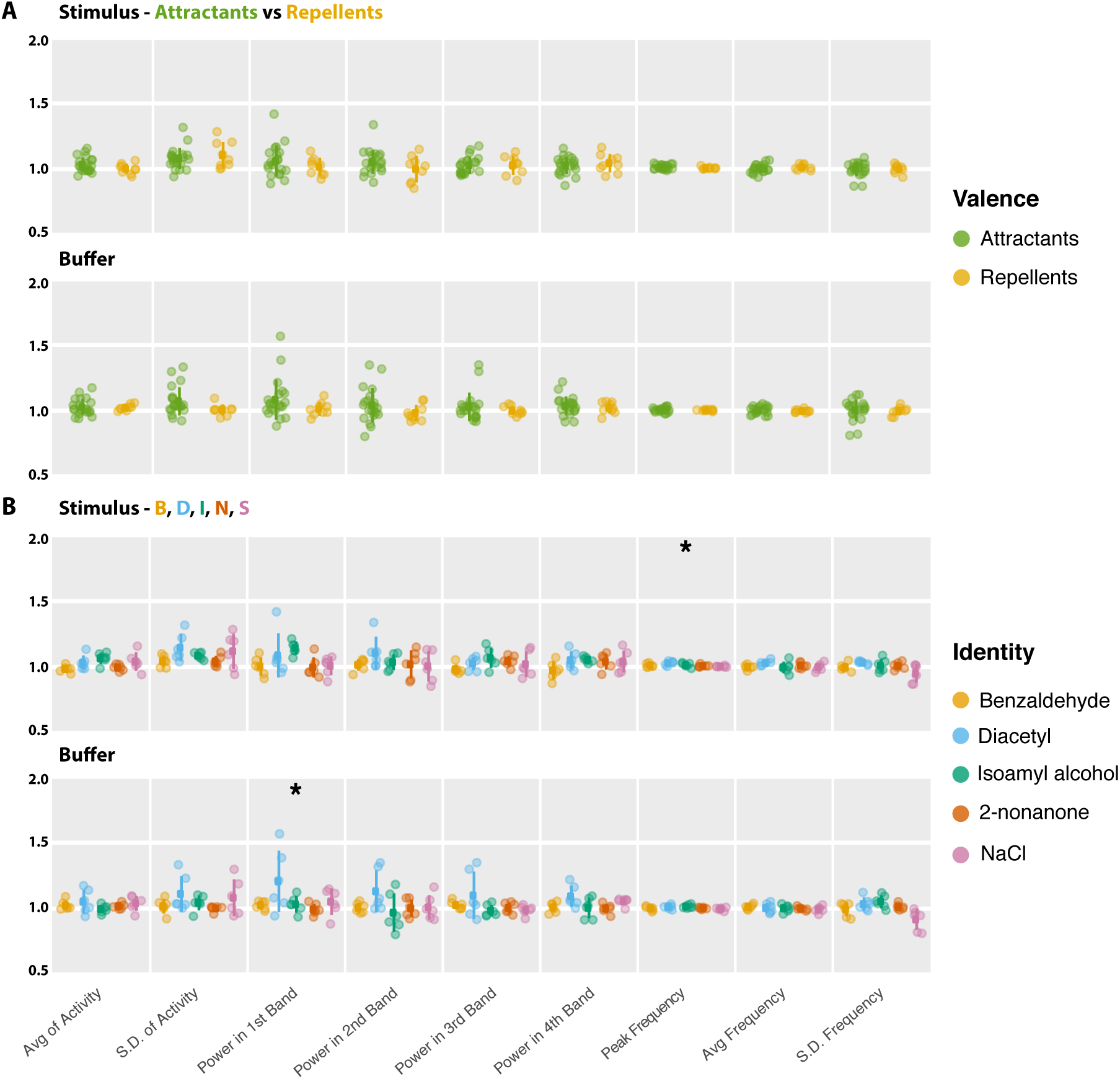
Stimulus valence and identity have distinct effects on neural activity on stimulus offset. Repellents effect no change in neural activity on stimulus offset (A). On the other hand, stimulus identity affects the peak frequency of neural activity on stimulus offset (B). Buffer offset affected average power in the 1st frequency band (B). Avg and S.D. of activity refer to average and standard deviation of neural activity. Power in 1st, 2nd, 3rd, and 4th bands refer to average power in the frequency ranges from 0.07 - 0.2 Hz, 0.2 - 0.34 Hz, 0.34 - 0.47 Hz, and 0.47 - 0.6 Hz. Peak frequency is the frequency with the most power in a 30-second bin, and avg frequency and S.D. frequency are the average and standard deviation, respectively, of the frequencies with the most power in a sliding-window bin covering a 30-second period. N = 21 for attractants and N = 9 for repellents (A), and N = 6 for each chemical stimulus (B). * p < 0.025, *** p < 0.0005, by likelihood ratio test on full and null generalized linear mixed-effects model, where the former included either valence or identity as a fixed effect.

**Supplemental Figure S4.**
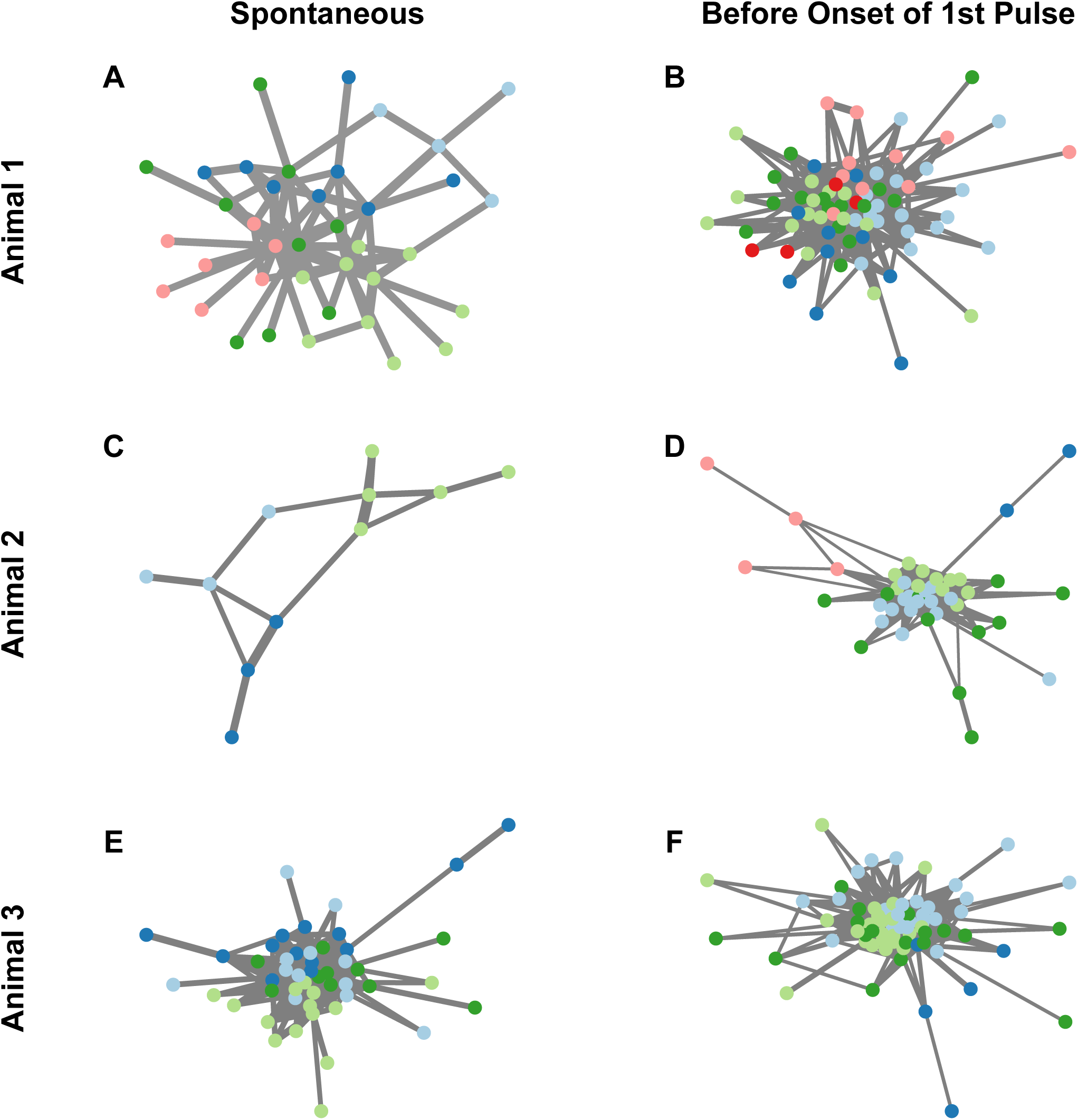
The neural networks of worms vary both within and across worms. The networks depicted here, for three example worms, show neurons (circles) connected by lines (edges); the thickness of the edges correspond to the amount of normalized mutual information between the two linked neurons. All networks are based on neural activity observed in the 30-second period between 30 seconds and 1 minute into the beginning of either the Spontaneous (A, C, E) or Stimulus (B, D, F) session for three worms. Neurons with the same color belong to the same module. Some worms have different numbers of modules in the absence of stimulus (A has 5 modules, B has 6 modules), while others have different numbers of strongly interacting neurons (C, D), and still others look fairly similar (E, F). All edges less than 0.2 were removed for the purpose of visualization.

**Supplemental Figure S5.**
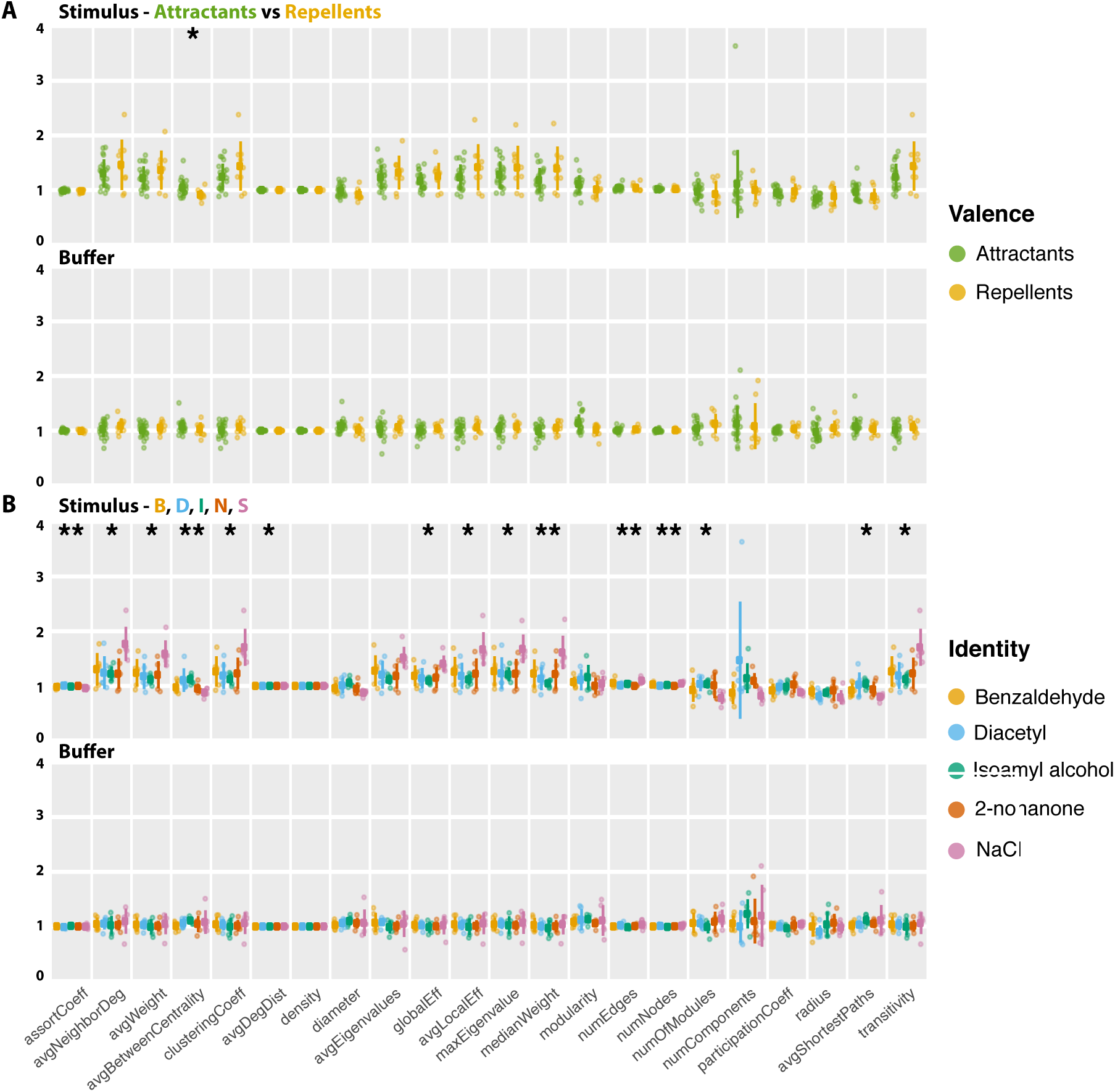
Stimulus valence and identity modulate distinct network features on stimulus onset. All 22 graph-theoretic features on the x-axes are plotted according to stimulus valence (A) and identity (B), and depict results from networks computed on stimulus onset during Buffer and Stimulus sessions. Network features are derived from adjacency matrices constructed with normalized mutual information. Colored dots indicate the mean across 7 pulses. N = 21 for attractants and N = 9 for repellents (A), and N = 6 for each chemical stimulus (B). * p < 0.025, ** p < 0.005, by likelihood ratio test on full and null generalized linear mixed-effects model, where the former included either valence or identity as a fixed effect.

**Supplemental Figure S6.**
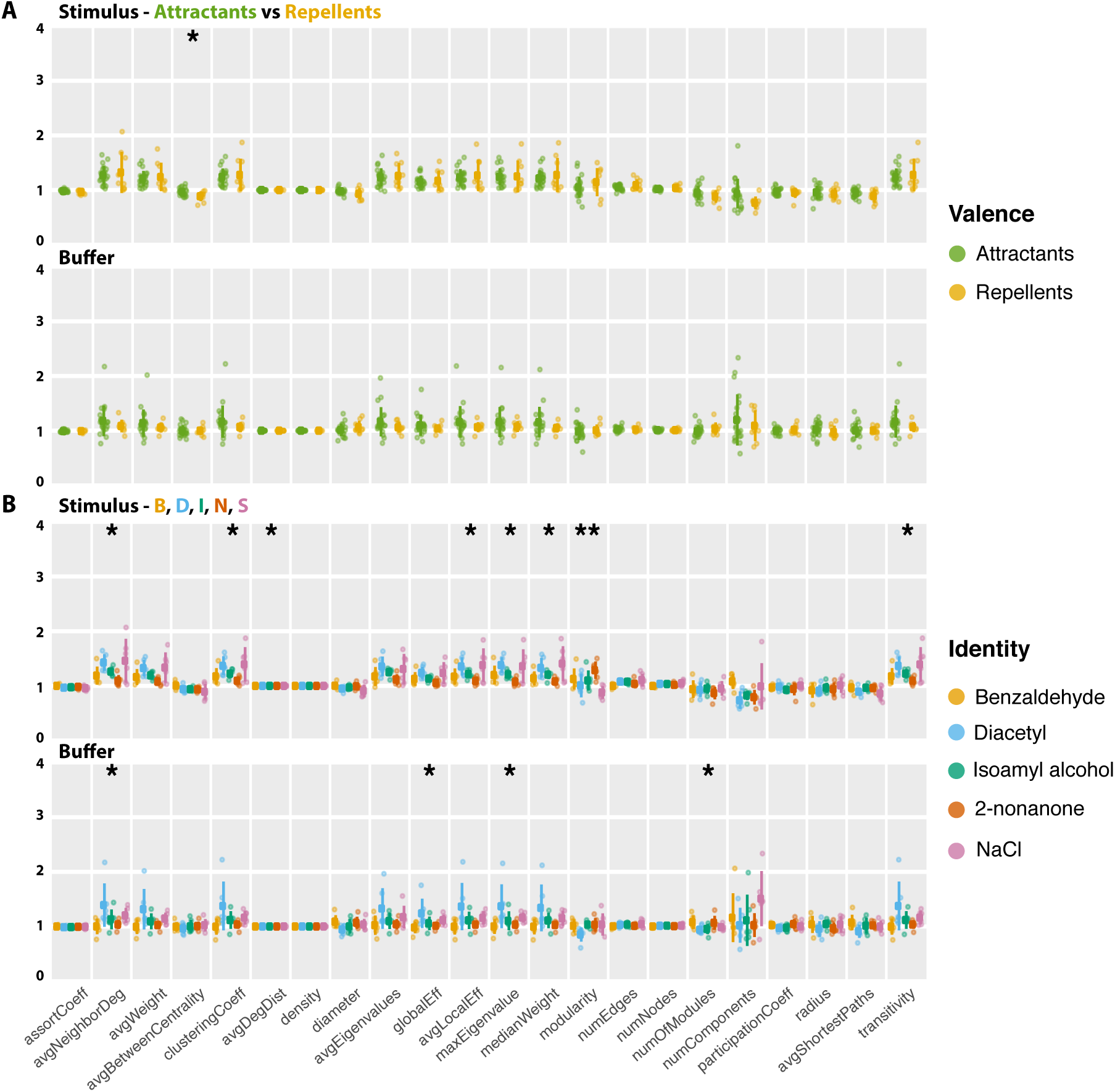
Stimulus valence and identity modulate distinct network features on stimulus offset. All 22 graph-theoretic features on the x-axes are plotted according to stimulus valence (A) and identity (B), and depict results from networks computed on stimulus offset during Buffer and Stimulus sessions. Network features are derived from adjacency matrices constructed with normalized mutual information. Colored dots indicate the mean across 7 pulses. N = 21 for attractants and N = 9 for repellents (A), and N = 6 for each chemical stimulus (B). * p < 0.025, ** p < 0.005, by likelihood ratio test on full and null generalized linear mixed-effects model, where the former included either valence or identity as a fixed effect.

**Supplemental Figure S7.**
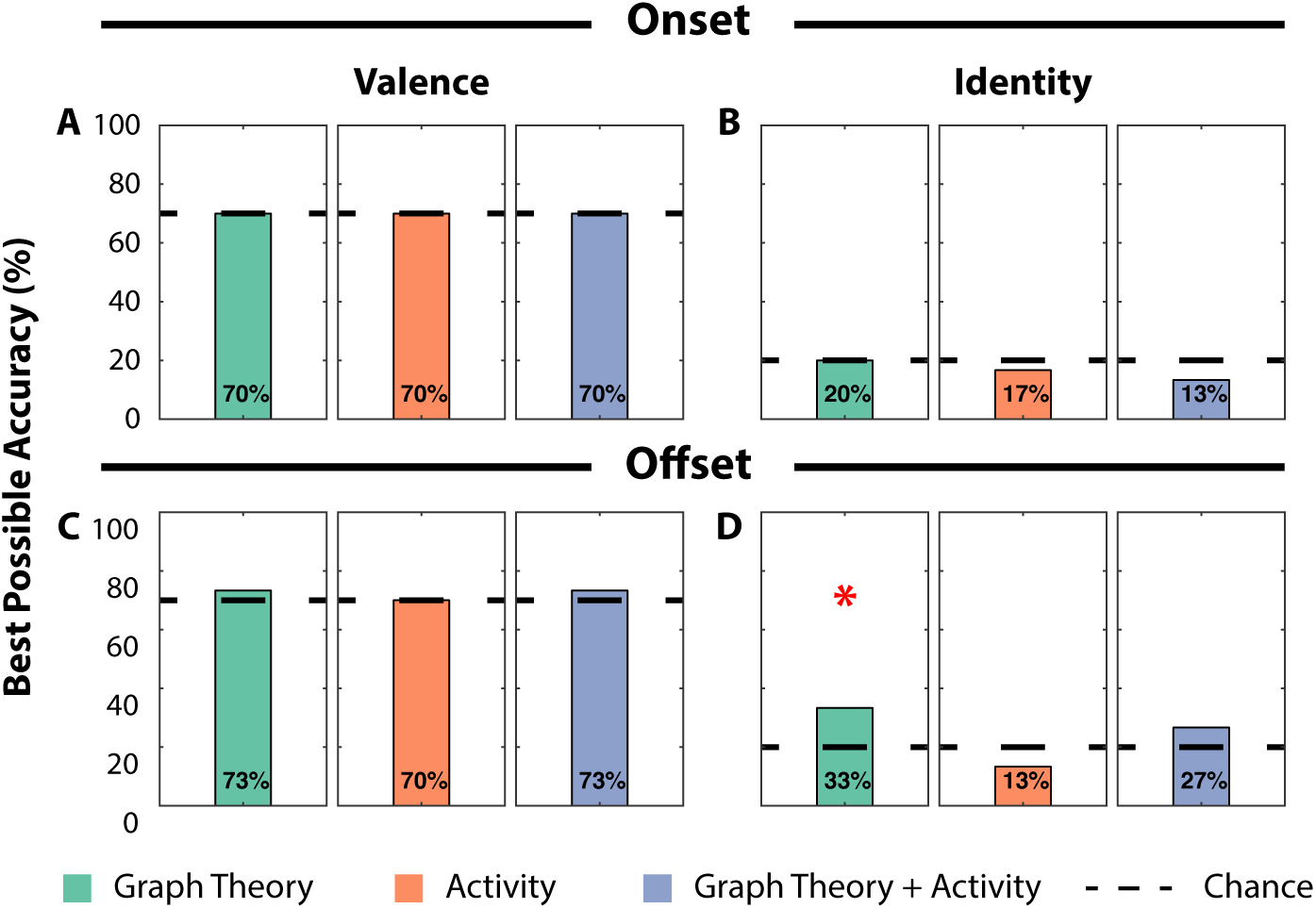
Logistic regression classifier attains above-chance classification accuracy for stimulus identity on Buffer sessions. Classification accuracy achieved on the first pulse of Buffer sessions using a Logistic Regression classifier trained using network features (green), neural activity features (orange) or both (blue). Accuracies are shown for stimulus valence and identity at stimulus onset (A, B) and offset (C, D), respectively. Red asterisks indicate significantly above-chance classification accuracy as determined with a permutation test (*p < 0.025). Black dotted lines indicate theoretical chance accuracy.

**Supplemental Figure S8.**
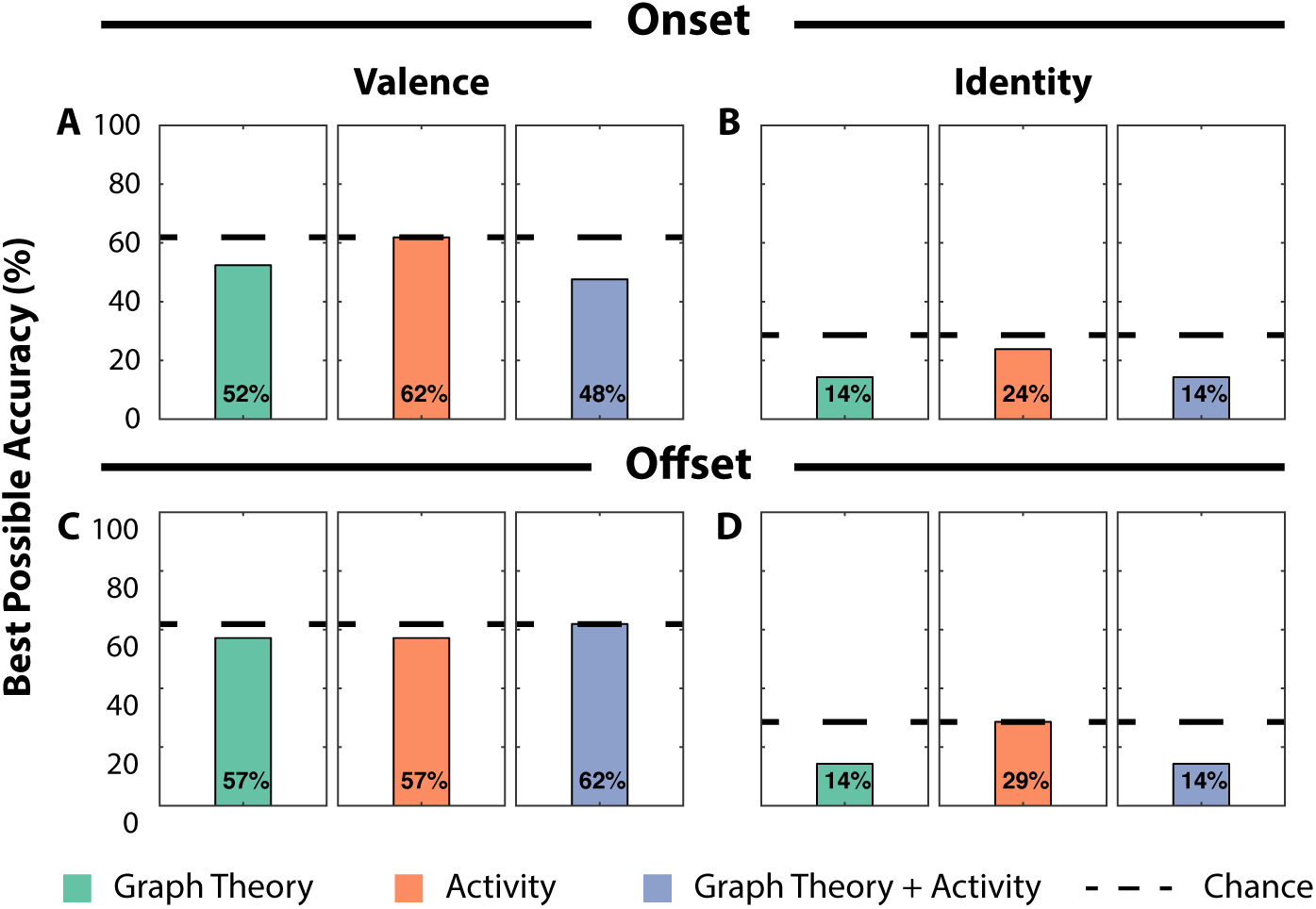
Logistic regression classifier does not attain above-chance classification accuracy for stimulus identity on Buffer sessions in‘naïve’ animals. Classification accuracy achieved on the first pulse of Buffer sessions in naïve animals using a Logistic Regression classifier trained using network features (green), neural activity features (orange) or both (blue). Accuracies are shown for stimulus valence and identity at stimulus onset (A, B) and offset (C, D), respectively. We do not achieve significantly above-chance classification accuracy, assessed with permutation testing (p > 0.05). Black dotted lines indicate theoretical chance accuracy.

**Table S1.** Definition of graph-theoretic features studied in this manuscript. The twenty-two listed features are grouped into one of five classes: basic structure (of the adjacency matrix), functional segregation (measures of the decomposability of the network), functional integration (the potential for disparate parts of the network to communicate), centrality (the importance of any one neuron to network communication), and resilience to perturbations, such as lesions (measures of how robust the system is to disruptions at individual nodes). For a review, see (1).

| Graph Theory Feature | Description |
| --- | --- |
| *Basic structure* |  |
| numNodes | the number of neurons that had at least one non-zero weight to another neuron |
| numEdges | the number of edges |
| density | the density of edges (i.e., the number of actual edges divided by the total possible number of edges in a fully-connected network) |
| numComponents | the number of isolated subgraphs in the network |
| avgWeight | the average weight of the network |
| medWeight | the median weight of the network |
| avgEigenvalue | the mean of the positive eigenvalues of the adjacency matrix |
| maxEigenvalue | the largest eigenvalue of the adjacency matrix |
| *Functional segregation* |  |
| avgClusteringCoeff | the average over all neurons of the fraction of 3-neuron clusters, or triangles, around each neuron |
| transitivity | a globally normalized version of the clustering coefficient |
| avgLocalEff | the mean local efficiency (i.e., the average over all neurons of the lengths of the shortest paths between two of the neuron's neighbors) |
| modularity | the extent to which a network can be divided into clusters, or modules, of neurons with dense connections amongst themselves, and sparse connections to neurons in other clusters |
| numModules | the number of modules |
| *Functional integration* |  |
| avgShortestPath | the average shortest path between all pairs of neurons in the network |
| globalEff | the global efficiency, or the average inverse shortest path length |
| radius | the smallest shortest path connecting any two neurons |
| diameter | the largest shortest path connecting any two neurons |
| *Centrality* |  |
| avgParticipationCoeff | the degree to which a neuron communicates with neurons in different modules |
| avgBetweennessCentrality | the average of the fraction of shortest paths linking any two neurons that pass through a given neuron |
| *Resilience to perturbations* |  |
| avgDegDist | the average of each neuron's degree, or sum of the edge weights to other neurons |
| assortCoeff | the correlation coefficient between the degrees of two connected neurons, where a negative number indicates neurons with a high degree are connected to neurons with a low degree |
| avgNeighborDegree | the average degree of the neighbors of a given node, averaged over all nodes |

**Table S2.** Graph-theoretic results from likelikood ratio test on generalized linear mixed-effects models. Results from the likelihood ratio test applied on a full vs null model. The full model includes information on either Valence or Identity in addition to the null model. The null model includes information on animal ID and time since first pulse. The p-values in red indicate a significant difference in the data’s likelihood when explained with the full model vs the null model; hence, the parameter (i.e., Valence or Identity) significantly improved model fit.

| Session | Pulse Switch | Graph Theory Feature | Valence | Identity |
| --- | --- | --- | --- | --- |
| Buffer | Onset | assortCoeff | 0.349 | 0.550 |
|  |  | avgBetweenCentrality | 0.288 | 0.665 |
|  |  | avgClusteringCoeff | 0.335 | 0.939 |
|  |  | avgDegDist | 0.448 | 0.688 |
|  |  | avgEigenvalue | 0.520 | 0.768 |
|  |  | avgLocalEff | 0.318 | 0.939 |
|  |  | avgNeighborDeg | 0.297 | 0.915 |
|  |  | avgParticipationCoeff | 0.235 | 0.362 |
|  |  | avgShortestPath | 0.329 | 0.607 |
|  |  | avgWeight | 0.341 | 0.952 |
|  |  | density | 0.530 | 0.649 |
|  |  | diameter | 0.166 | 0.872 |
|  |  | globalEff | 0.311 | 0.955 |
|  |  | maxEigenvalue | 0.405 | 0.970 |
|  |  | medWeight | 0.219 | 0.912 |
|  |  | modularity | 0.034 | 0.918 |
|  |  | numComponents | 0.626 | 0.756 |
|  |  | numEdges | 0.297 | 0.270 |
|  |  | numModules | 0.152 | 0.457 |
|  |  | numNodes | 0.303 | 0.367 |
|  |  | radius | 0.246 | 0.432 |
|  |  | transitivity | 0.335 | 0.939 |
|  | Offset | assortCoeff | 0.930 | 0.904 |
|  |  | avgBetweenCentrality | 0.940 | 0.866 |
|  |  | avgClusteringCoeff | 0.307 | 0.032 |
|  |  | avgDegDist | 0.979 | 0.840 |
|  |  | avgEigenvalue | 0.261 | 0.098 |
|  |  | avgLocalEff | 0.306 | 0.032 |
|  |  | avgNeighborDeg | 0.322 | **0.011** |
|  |  | avgParticipationCoeff | 0.508 | 0.241 |
|  |  | avgShortestPath | 0.982 | 0.165 |
|  |  | avgWeight | 0.334 | 0.031 |
|  |  | density |  | 0.138 |
|  |  | diameter | 0.239 | 0.146 |
|  |  | globalEff | 0.386 | 0.022 |
|  |  | maxEigenvalue | 0.370 | 0.015 |
|  |  | medWeight | 0.250 | 0.049 |
|  |  | modularity | 0.706 | 0.048 |
|  |  | numComponents | 0.606 | 0.222 |
|  |  | numEdges | 0.695 | 0.484 |
|  |  | numModules | 0.311 | 0.084 |
|  |  | numNodes | 0.819 | 0.658 |
|  |  | radius | 0.398 | 0.676 |
|  |  | transitivity | 0.307 | 0.032 |
| Stimulus | Onset | assortCoeff | 0.486 | 0.002 |
|  |  | avgBetweenCentrality | 0.013 | 0.003 |
|  |  | avgClusteringCoeff | 0.154 | 0.005 |
|  |  | avgDegDist | 0.808 | 0.008 |
|  |  | avgEigenvalue | 0.388 | 0.028 |
|  |  | avgLocalEff | 0.158 | 0.006 |
|  |  | avgNeighborDeg | 0.285 | 0.008 |
|  |  | avgParticipationCoeff | 0.583 | 0.095 |
|  |  | avgShortestPath | 0.153 | 0.016 |
|  |  | avgWeight | 0.195 | 0.009 |
|  |  | density | 0.865 | 0.089 |
|  |  | diameter | 0.076 | 0.028 |
|  |  | globalEff | 0.259 | 0.020 |
|  |  | maxEigenvalue | 0.322 | 0.016 |
|  |  | medWeight | 0.079 | 0.003 |
|  |  | modularity | 0.141 | 0.495 |
|  |  | numComponents | 0.766 | 0.032 |
|  |  | numEdges | 0.778 | 0.003 |
|  |  | numModules | 0.481 | 0.021 |
|  |  | numNodes | 0.733 | 0.003 |
|  |  | radius | 0.405 | 0.242 |
|  |  | transitivity | 0.154 | 0.005 |
|  | Offset | assortCoeff | 0.075 | 0.063 |
|  |  | avgBetweenCentrality | 0.021 | 0.267 |
|  |  | avgClusteringCoeff | 0.654 | 0.020 |
|  |  | avgDegDist | 0.164 | 0.064 |
|  |  | avgEigenvalue | 0.692 | 0.126 |
|  |  | avgLocalEff | 0.649 | 0.022 |
|  |  | avgNeighborDeg | 0.638 | 0.012 |
|  |  | avgParticipationCoeff | 0.410 | 0.325 |
|  |  | avgShortestPath | 0.158 | 0.196 |
|  |  | avgWeight | 0.696 | 0.029 |
|  |  | density | 0.469 | 0.232 |
|  |  | diameter | 0.131 | 0.247 |
|  |  | globalEff | 0.729 | 0.050 |
|  |  | maxEigenvalue | 0.938 | 0.017 |
|  |  | medWeight | 0.471 | 0.011 |
|  |  | modularity | 0.304 | 0.002 |
|  |  | numComponents | 0.056 | 0.041 |
|  |  | numEdges | 0.113 | 0.030 |
|  |  | numModules | 0.176 | 0.706 |
|  |  | numNodes | 0.110 | 0.030 |
|  |  | radius | 0.587 | 0.686 |
|  |  | transitivity | 0.654 | 0.020 |

**Table S3.**
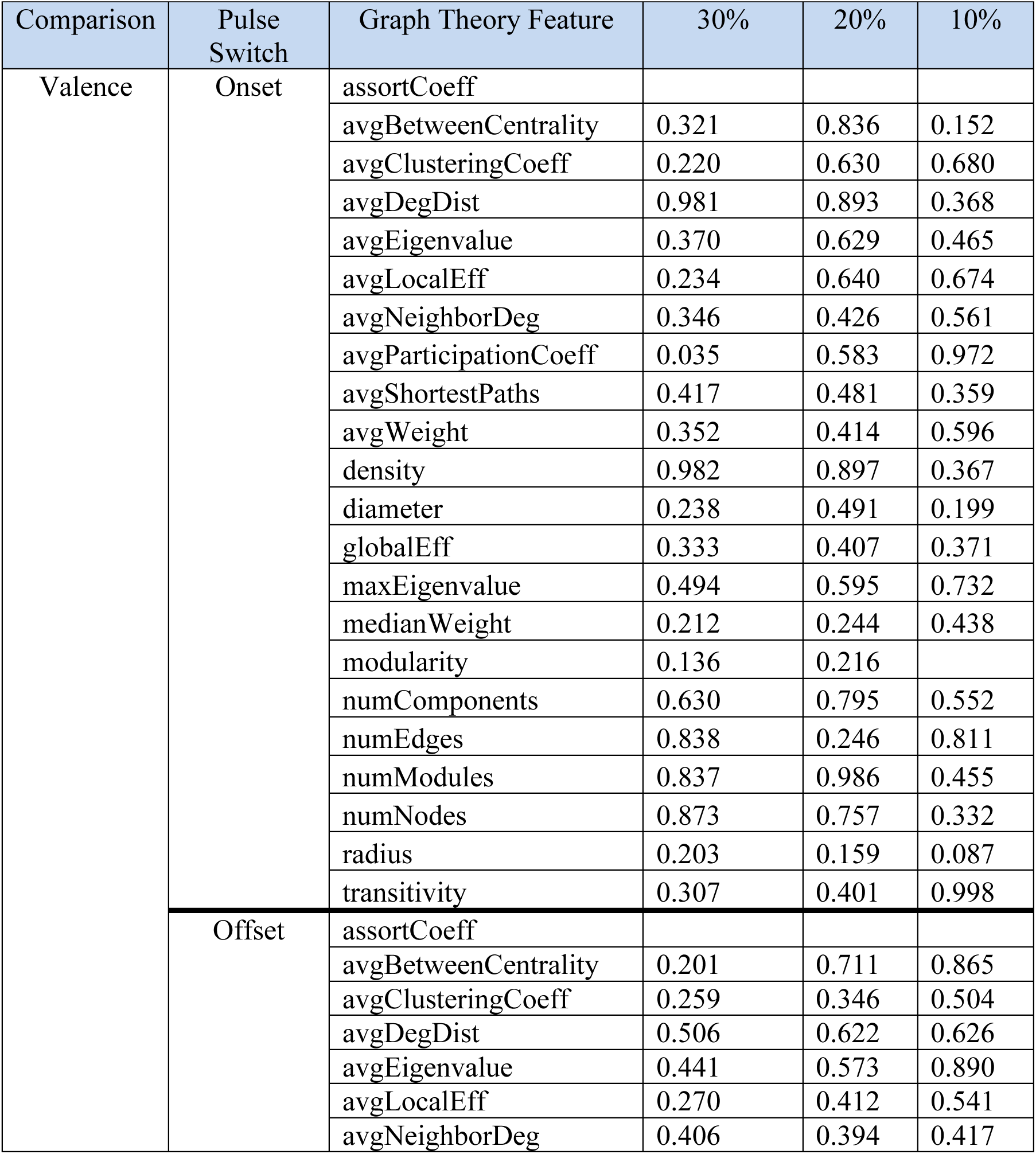
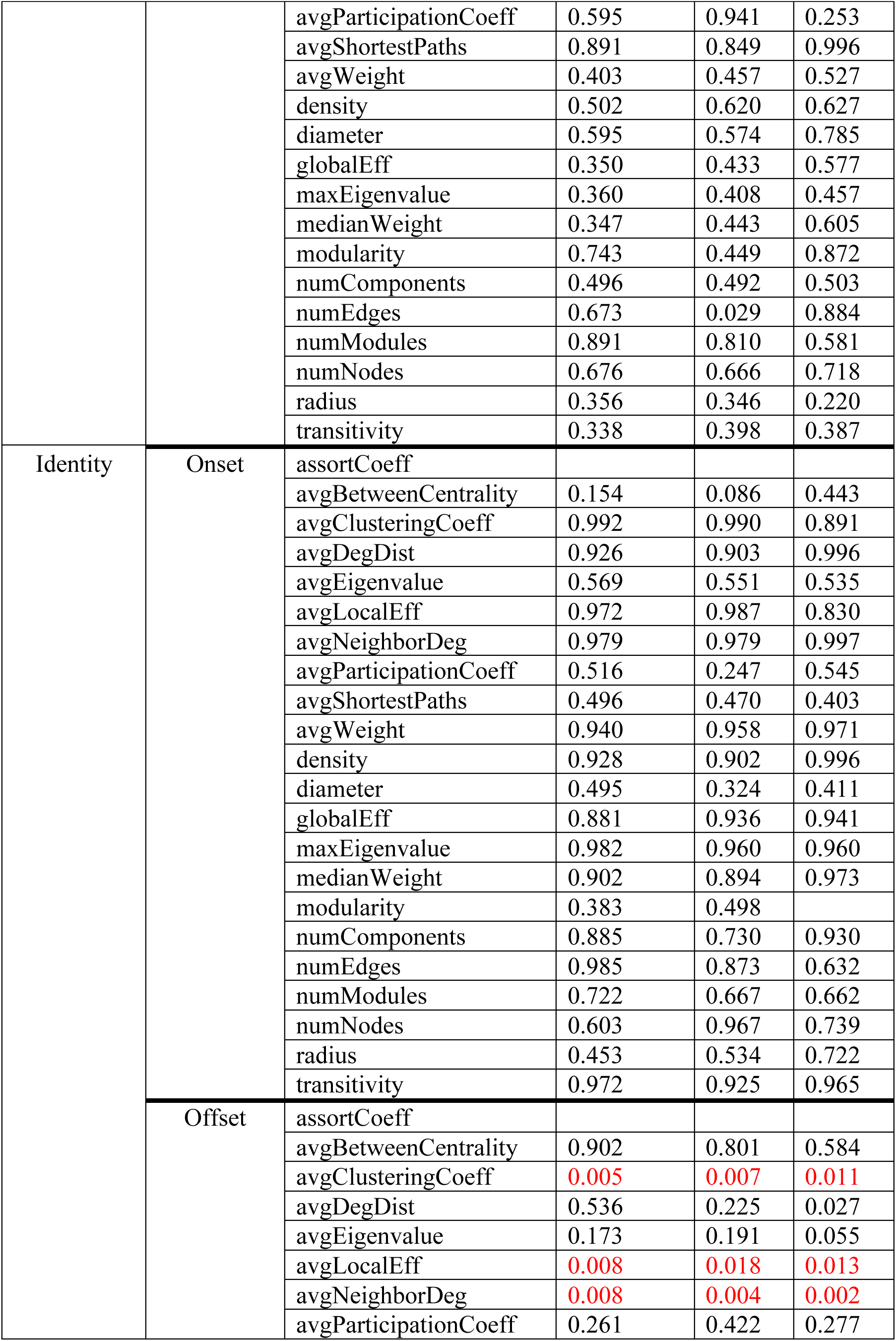
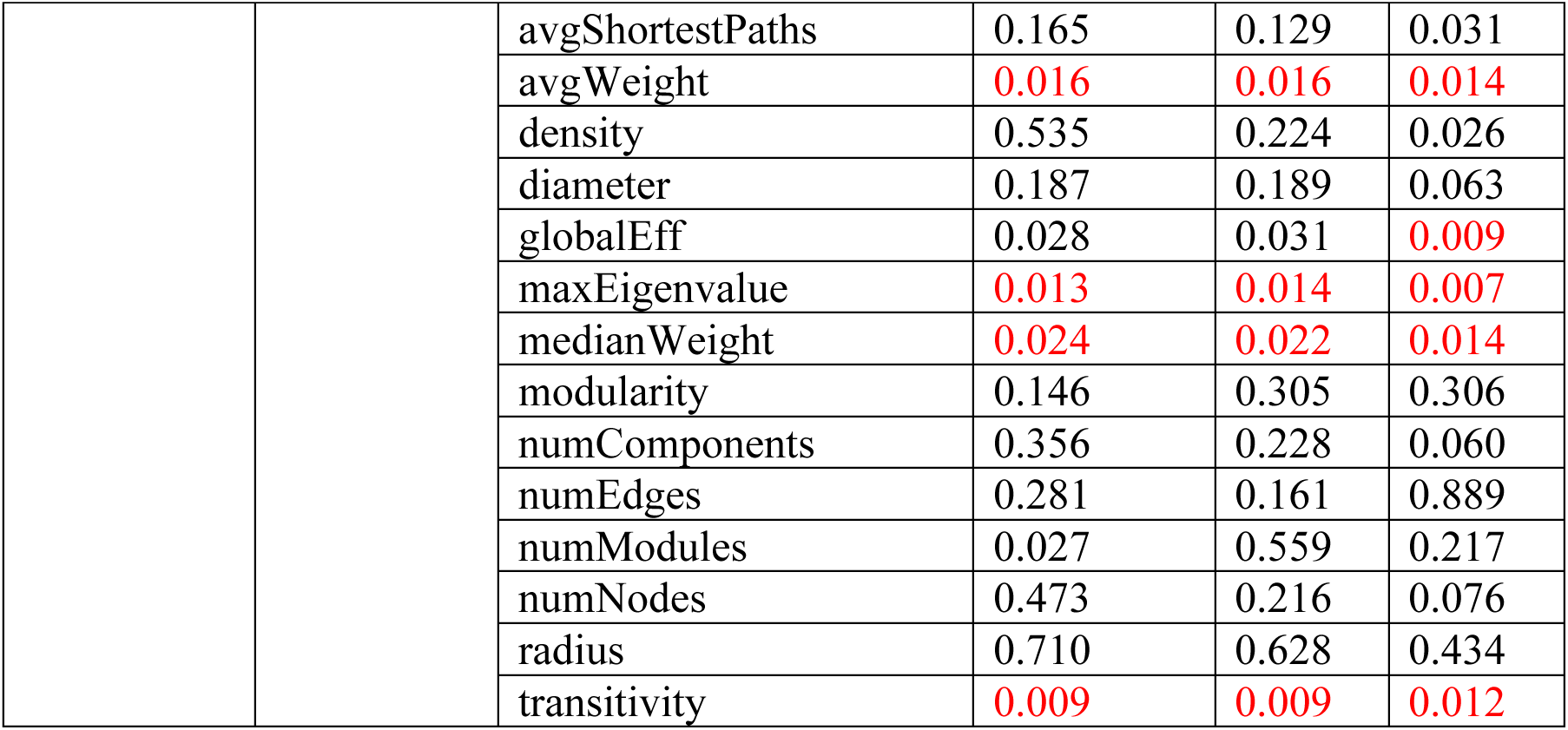
Graph-theoretic results from likelikood ratio test on generalized linear mixed-effects models for the Buffer session on thresholded networks. Only the top 30%, 20%, or 10% of weights were kept in each network (columns). Results from the likelihood ratio test applied on a full vs null model. The full model includes information on either Valence or Identity in addition to the null model. The null model includes information on animal ID and time since first pulse. The p-values in red indicate a significant difference in the data’s likelihood when explained with the full model vs the null model; hence, the parameter (i.e., Valence or Identity) significantly improved model fit. The ‘assortCoeff’ cells are empty because they had negative numbers which could not be fit by a gamma distribution GLME. Other cells are empty because extreme outliers prevented convergence.

**Table S4.** Graph-theoretic results from likelikood ratio test on generalized linear mixed-effects models for the Stimulus session on thresholded networks. Only the top 30%, 20%, or 10% of weights were kept in each network (columns). Results from the likelihood ratio test applied on a full vs null model. The full model includes information on either Valence or Identity in addition to the null model. The null model includes information on animal ID and time since first pulse. The p-values in red indicate a significant difference in the data’s likelihood when explained with the full model vs the null model; hence, the parameter (i.e., Valence or Identity) significantly improved model fit. The ‘assortCoeff’ cells are empty because they had negative numbers which could not be fit by a gamma distribution GLME. Other cells are empty because extreme outliers prevented convergence.

| Comparison | Pulse Switch | Graph Theory Feature | 30% | 20% | 10% |
| --- | --- | --- | --- | --- | --- |
| Valence | Onset | assortCoeff |  |  |  |
|  |  | avgBetweenCentrality | 0.009 | 0.114 | 0.814 |
|  |  | avgClusteringCoeff | 0.275 | 0.400 | 0.582 |
|  |  | avgDegDist | 0.631 | 0.978 | 0.954 |
|  |  | avgEigenvalue | 0.308 | 0.533 | 0.714 |
|  |  | avgLocalEff | 0.211 | 0.346 | 0.618 |
|  |  | avgNeighborDeg | 0.333 | 0.439 | 0.708 |
|  |  | avgParticipationCoeff | 0.495 | 0.474 | 0.635 |
|  |  | avgShortestPaths | 0.159 | 0.216 | 0.327 |
|  |  | avgWeight | 0.271 | 0.328 | 0.436 |
|  |  | density | 0.631 | 0.974 | 0.969 |
|  |  | diameter | 0.101 | 0.224 | 0.185 |
|  |  | globalEff | 0.318 | 0.438 | 0.610 |
|  |  | maxEigenvalue | 0.484 | 0.648 | 0.790 |
|  |  | medianWeight | 0.195 | 0.252 | 0.340 |
|  |  | modularity | 0.578 | 0.615 | 0.738 |
|  |  | numComponents | 0.106 | 0.526 | 0.556 |
|  |  | numEdges | 0.486 | 0.685 | 0.207 |
|  |  | numModules | 0.342 | 0.534 | 0.669 |
|  |  | numNodes | 0.536 | 0.824 | 0.601 |
|  |  | radius | 0.247 | 0.321 | 0.370 |
|  |  | transitivity | 0.397 | 0.545 | 0.633 |
|  | Offset | assortCoeff |  |  |  |
|  |  | avgBetweenCentrality | 0.001 | 0.000 | 0.731 |
|  |  | avgClusteringCoeff | 0.913 | 0.905 | 0.352 |
|  |  | avgDegDist | 0.155 | 0.554 | 0.183 |
|  |  | avgEigenvalue | 0.606 | 0.784 | 0.787 |
|  |  | avgLocalEff | 0.588 | 0.862 | 0.370 |
|  |  | avgNeighborDeg | 0.996 | 0.926 | 0.456 |
|  |  | avgParticipationCoeff | 0.407 | 0.746 | 0.684 |
|  |  | avgShortestPaths | 0.132 | 0.096 | 0.294 |
|  |  | avgWeight | 0.630 | 0.772 | 0.940 |
|  |  | density | 0.155 | 0.552 | 0.180 |
|  |  | diameter | 0.007 | 0.003 | 0.172 |
|  |  | globalEff | 0.840 | 0.833 | 0.606 |
|  |  | maxEigenvalue | 0.561 | 0.536 | 0.285 |
|  |  | medianWeight | 0.468 | 0.622 |  |
|  |  | modularity | 0.188 | 0.281 | 0.277 |
|  |  | numComponents | 0.467 | 0.089 | 0.131 |
|  |  | numEdges | 0.487 | 0.264 | 0.688 |
|  |  | numModules | 0.236 | 0.179 | 0.851 |
|  |  | numNodes | 0.224 | 0.842 | 0.373 |
|  |  | radius | 0.717 | 0.918 | 0.332 |
|  |  | transitivity | 0.768 | 0.552 | 0.297 |
| Identity | Onset | assortCoeff |  |  |  |
|  |  | avgBetweenCentrality | 0.011 | 0.422 | 0.223 |
|  |  | avgClusteringCoeff | 0.009 | 0.025 | 0.082 |
|  |  | avgDegDist | 0.095 | 0.574 | 0.870 |
|  |  | avgEigenvalue | 0.017 | 0.042 | 0.206 |
|  |  | avgLocalEff | 0.008 | 0.034 | 0.089 |
|  |  | avgNeighborDeg | 0.019 | 0.034 | 0.080 |
|  |  | avgParticipationCoeff | 0.045 | 0.000 | 0.003 |
|  |  | avgShortestPaths | 0.018 | 0.041 | 0.116 |
|  |  | avgWeight | 0.008 | 0.013 | 0.022 |
|  |  | density | 0.095 | 0.576 | 0.870 |
|  |  | diameter | 0.109 | 0.187 | 0.554 |
|  |  | globalEff | 0.025 | 0.063 | 0.183 |
|  |  | maxEigenvalue | 0.039 | 0.088 | 0.204 |
|  |  | medianWeight | 0.005 | 0.011 | 0.021 |
|  |  | modularity | 0.584 | 0.614 | 0.575 |
|  |  | numComponents | 0.018 | 0.240 | 0.650 |
|  |  | numEdges | 0.096 | 0.699 | 0.467 |
|  |  | numModules | 0.051 | 0.106 | 0.505 |
|  |  | numNodes | 0.047 | 0.387 | 0.722 |
|  |  | radius | 0.162 | 0.171 | 0.312 |
|  |  | transitivity | 0.012 | 0.022 | 0.073 |
|  | Offset | assortCoeff |  |  |  |
|  |  | avgBetweenCentrality | 0.002 | 0.013 | 0.420 |
|  |  | avgClusteringCoeff | 0.009 | 0.011 | 0.004 |
|  |  | avgDegDist | 0.440 | 0.325 | 0.135 |
|  |  | avgEigenvalue | 0.256 | 0.062 | 0.098 |
|  |  | avgLocalEff | 0.017 | 0.019 | 0.006 |
|  |  | avgNeighborDeg | 0.008 | 0.010 | 0.005 |
|  |  | avgParticipationCoeff | 0.271 | 0.070 | 0.114 |
|  |  | avgShortestPaths | 0.160 | 0.098 | 0.072 |
|  |  | avgWeight | 0.019 | 0.023 | 0.035 |
|  |  | density | 0.439 | 0.323 | 0.134 |
|  |  | diameter | 0.129 | 0.040 | 0.067 |
|  |  | globalEff | 0.037 | 0.033 | 0.035 |
|  |  | maxEigenvalue | 0.009 | 0.008 | 0.009 |
|  |  | medianWeight | 0.012 | 0.016 | 0.029 |
|  |  | modularity | 0.118 | 0.089 | 0.609 |
|  |  | numComponents | 0.833 | 0.227 | 0.095 |
|  |  | numEdges | 0.829 | 0.248 | 0.620 |
|  |  | numModules | 0.824 | 0.698 | 0.031 |
|  |  | numNodes | 0.394 | 0.615 | 0.324 |
|  |  | radius | 0.512 | 0.291 | 0.141 |
|  |  | transitivity | 0.006 | 0.005 | 0.007 |

**Table S5.** Graph-theoretic results from likelikood ratio test on generalized linear mixed-effects models for the Buffer session on differently correlated networks. The positively (+) and negatively (-) correlated subnetworks are shown (columns). Results from the likelihood ratio test applied on a full vs null model. The full model includes information on either Valence or Identity in addition to the null model. The null model includes information on animal ID and time since first pulse. The p-values in red indicate a significant difference in the data’s likelihood when explained with the full model vs the null model; hence, the parameter (i.e., Valence or Identity) significantly improved model fit. The ‘assortCoeff’ cells are empty because they had negative numbers which could not be fit by a gamma distribution GLME. Other cells are empty because extreme outliers prevented convergence.

| Comparison | Pulse Switch | Graph Theory Feature | + | - |
| --- | --- | --- | --- | --- |
| Valence | Onset | assortCoeff |  |  |
|  |  | avgBetweenCentrality | 0.074 | 0.860 |
|  |  | avgClusteringCoeff | 0.490 |  |
|  |  | avgDegDist | 0.311 | 0.261 |
|  |  | avgEigenvalue | 0.419 | 0.364 |
|  |  | avgLocalEff | 0.394 | 0.532 |
|  |  | avgNeighborDeg | 0.193 | 0.308 |
|  |  | avgParticipationCoeff | 0.600 | 0.696 |
|  |  | avgShortestPaths | 0.234 | 0.340 |
|  |  | avgWeight | 0.392 | 0.385 |
|  |  | density | 0.321 | 0.258 |
|  |  | diameter | 0.626 | 0.698 |
|  |  | globalEff | 0.212 | 0.297 |
|  |  | maxEigenvalue | 0.511 | 0.715 |
|  |  | medianWeight | 0.137 | 0.387 |
|  |  | modularity | 0.048 | 0.857 |
|  |  | numComponents | 0.626 | 0.626 |
|  |  | numEdges | 0.113 | 0.705 |
|  |  | numModules | 0.527 | 0.815 |
|  |  | numNodes | 0.303 | 0.303 |
|  |  | radius | 0.397 | 0.639 |
|  |  | transitivity | 0.403 | **0.004** |
|  | Offset | assortCoeff |  |  |
|  |  | avgBetweenCentrality | 0.056 | 0.695 |
|  |  | avgClusteringCoeff | 0.344 | 0.205 |
|  |  | avgDegDist | 0.178 | 0.054 |
|  |  | avgEigenvalue | 0.344 | 0.345 |
|  |  | avgLocalEff | 0.319 | 0.214 |
|  |  | avgNeighborDeg | 0.284 | 0.395 |
|  |  | avgParticipationCoeff | 0.122 | 0.551 |
|  |  | avgShortestPaths | 0.531 | 0.512 |
|  |  | avgWeight | 0.371 | 0.322 |
|  |  | density | 0.180 | 0.052 |
|  |  | diameter | 0.648 | 0.443 |
|  |  | globalEff | 0.297 | 0.483 |
|  |  | maxEigenvalue | 0.439 | 0.566 |
|  |  | medianWeight | 0.269 | 0.251 |
|  |  | modularity | 0.071 | 0.331 |
|  |  | numComponents | 0.606 | 0.606 |
|  |  | numEdges | 0.238 | 0.388 |
|  |  | numModules | 0.997 | 0.113 |
|  |  | numNodes | 0.819 | 0.819 |
|  |  | radius | 0.535 | 0.243 |
|  |  | transitivity | 0.421 | 0.367 |
| Identity | Onset | assortCoeff |  |  |
|  |  | avgBetweenCentrality | 0.114 | 0.813 |
|  |  | avgClusteringCoeff | 0.972 | 0.757 |
|  |  | avgDegDist | 0.421 | 0.465 |
|  |  | avgEigenvalue | 0.759 | 0.748 |
|  |  | avgLocalEff | 0.952 | 0.124 |
|  |  | avgNeighborDeg | 0.654 | 0.914 |
|  |  | avgParticipationCoeff | 0.382 | 0.035 |
|  |  | avgShortestPaths | 0.560 | 0.637 |
|  |  | avgWeight | 0.933 | 0.902 |
|  |  | density | 0.436 | 0.452 |
|  |  | diameter | 0.813 | 0.508 |
|  |  | globalEff | 0.829 | 0.900 |
|  |  | maxEigenvalue | 0.698 | 0.844 |
|  |  | medianWeight | 0.903 | 0.801 |
|  |  | modularity | 0.087 | 0.779 |
|  |  | numComponents | 0.756 | 0.756 |
|  |  | numEdges | 0.099 | 0.978 |
|  |  | numModules | 0.787 | 0.165 |
|  |  | numNodes | 0.367 | 0.367 |
|  |  | radius | 0.447 | 0.576 |
|  |  | transitivity | 0.943 | 0.067 |
|  | Offset | assortCoeff |  |  |
|  |  | avgBetweenCentrality | 0.455 | 0.775 |
|  |  | avgClusteringCoeff | 0.049 | 0.022 |
|  |  | avgDegDist | 0.741 | 0.420 |
|  |  | avgEigenvalue | 0.038 | 0.008 |
|  |  | avgLocalEff | 0.040 | 0.006 |
|  |  | avgNeighborDeg | 0.039 | 0.010 |
|  |  | avgParticipationCoeff | 0.576 | 0.602 |
|  |  | avgShortestPaths | 0.136 | 0.123 |
|  |  | avgWeight | 0.036 | 0.029 |
|  |  | density | 0.746 | 0.416 |
|  |  | diameter | 0.233 | 0.190 |
|  |  | globalEff | 0.035 | 0.016 |
|  |  | maxEigenvalue | 0.048 | 0.011 |
|  |  | medianWeight | 0.052 | 0.033 |
|  |  | modularity | 0.472 | 0.862 |
|  |  | numComponents | 0.222 | 0.222 |
|  |  | numEdges | 0.419 | 0.543 |
|  |  | numModules | 0.379 | 0.231 |
|  |  | numNodes | 0.658 | 0.658 |
|  |  | radius | 0.637 | 0.171 |
|  |  | transitivity | 0.063 | 0.027 |

**Table S6.**
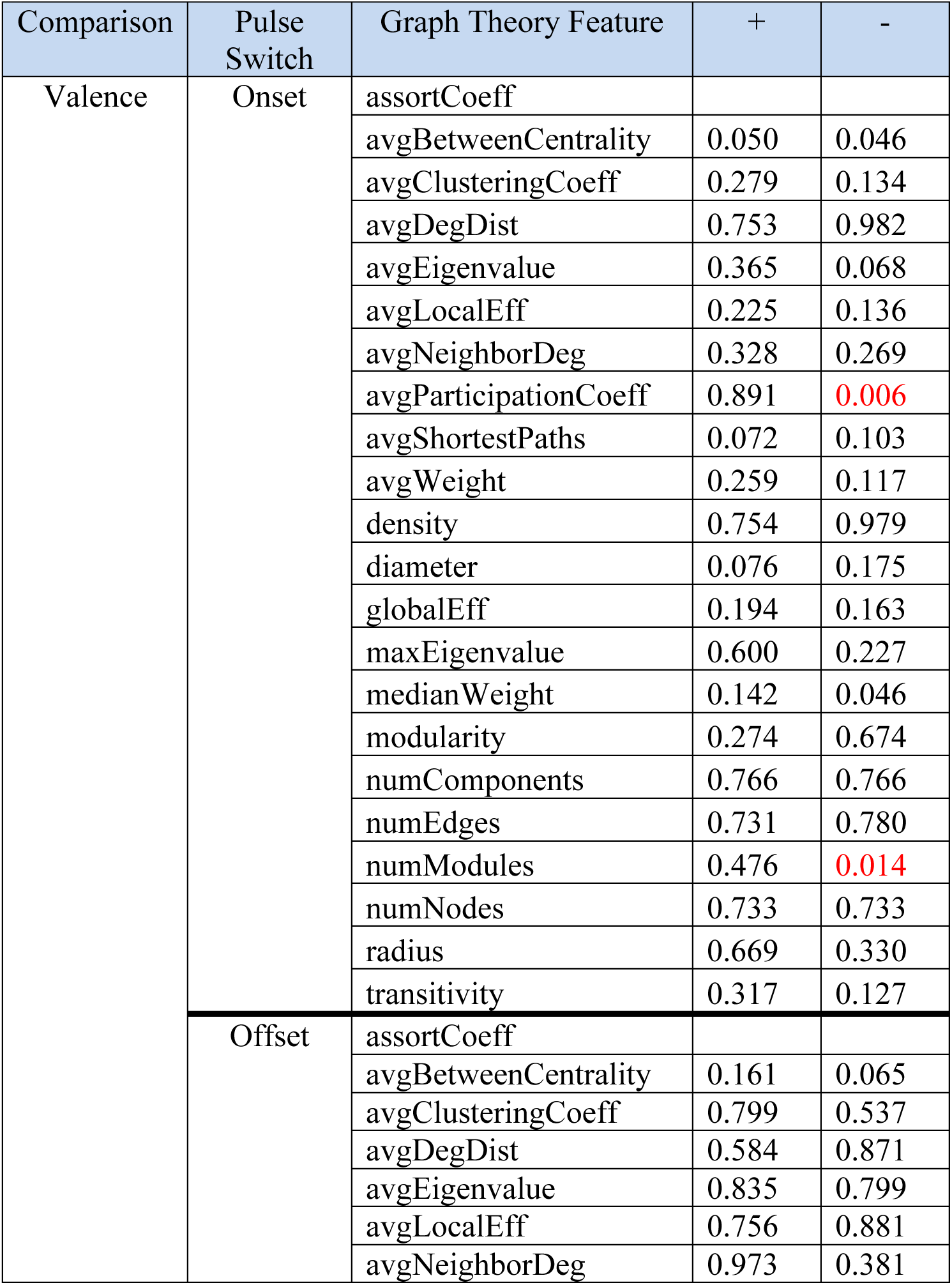
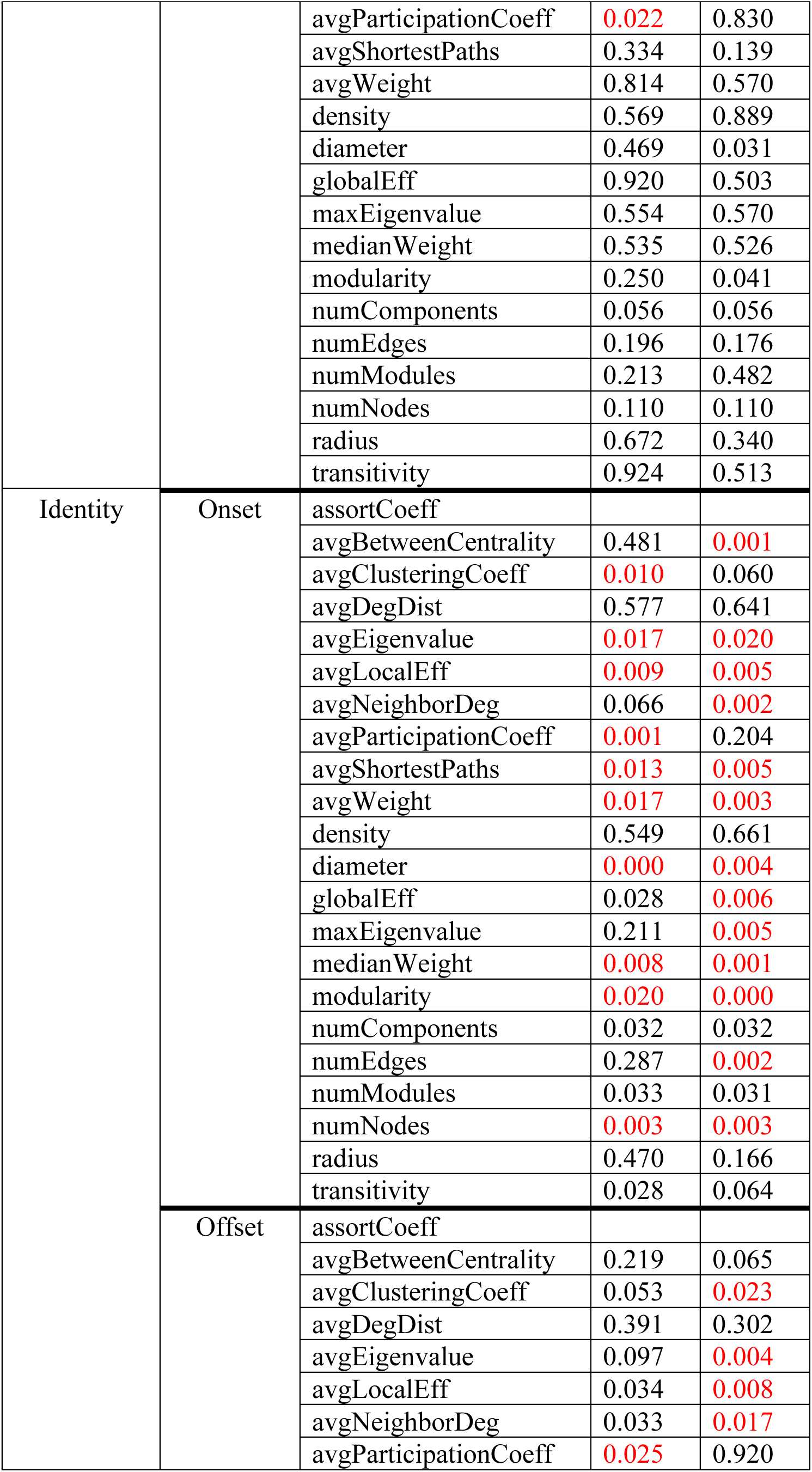
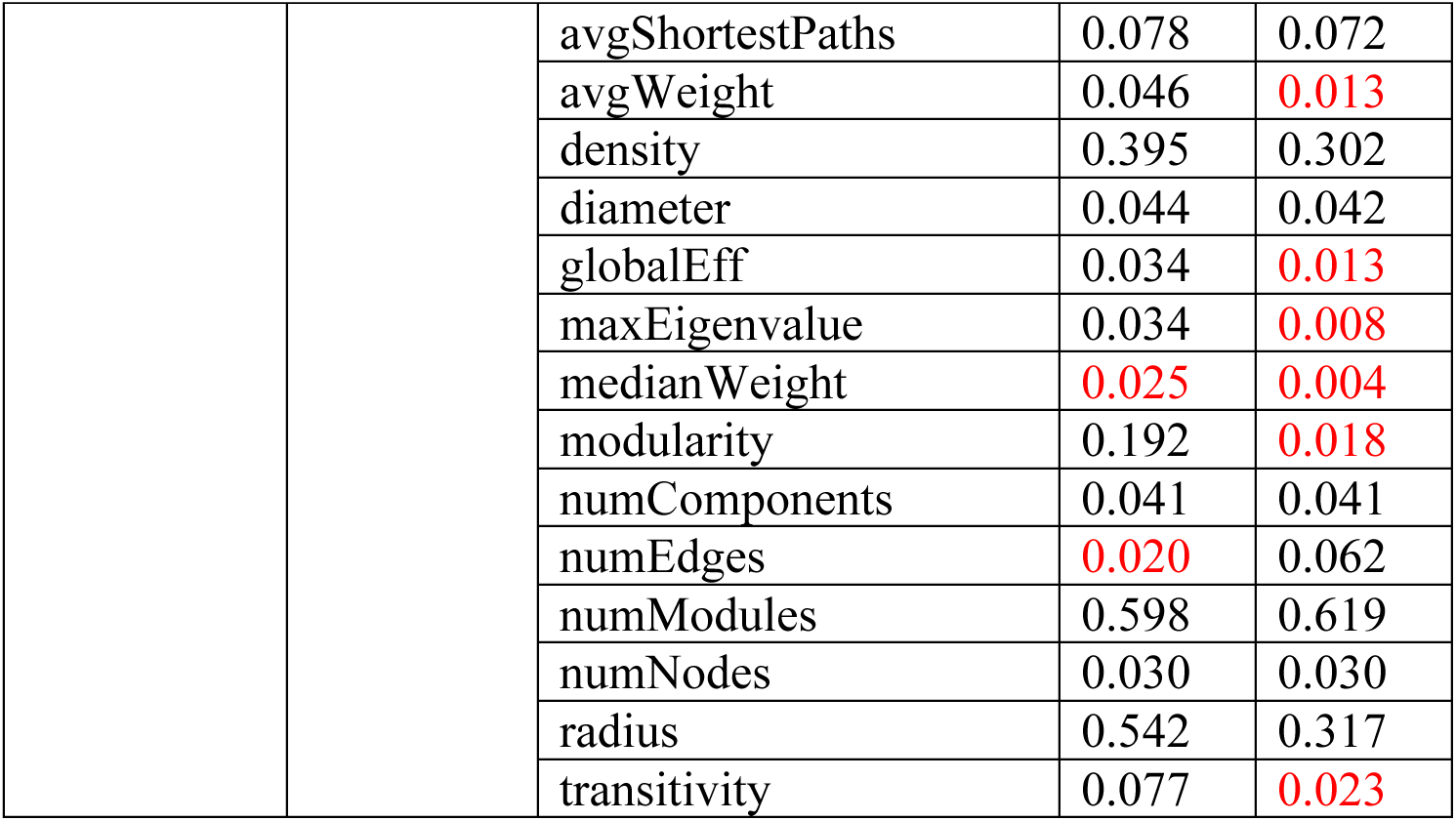
Graph-theoretic results from likelikood ratio test on generalized linear mixed-effects models for the Stimulus session on differently correlated networks. The positively (+) and negatively (-) correlated subnetworks are shown (columns). Results from the likelihood ratio test applied on a full vs null model. The full model includes information on either Valence or Identity in addition to the null model. The null model includes information on animal ID and time since first pulse. The p-values in red indicate a significant difference in the data’s likelihood when explained with the full model vs the null model; hence, the parameter (i.e., Valence or Identity) significantly improved model fit. The ‘assortCoeff’ cells are empty because they had negative numbers which could not be fit by a gamma distribution GLME. Other cells are empty because extreme outliers prevented convergence.

| Comparison | Pulse Switch | Graph Theory Feature | + | - |
| --- | --- | --- | --- | --- |
| Valence | Onset | assortCoeff |  |  |
|  |  | avgBetweenCentrality | 0.050 | 0.046 |
|  |  | avgClusteringCoeff | 0.279 | 0.134 |
|  |  | avgDegDist | 0.753 | 0.982 |
|  |  | avgEigenvalue | 0.365 | 0.068 |
|  |  | avgLocalEff | 0.225 | 0.136 |
|  |  | avgNeighborDeg | 0.328 | 0.269 |
|  |  | avgParticipationCoeff | 0.891 | 0.006 |
|  |  | avgShortestPaths | 0.072 | 0.103 |
|  |  | avgWeight | 0.259 | 0.117 |
|  |  | density | 0.754 | 0.979 |
|  |  | diameter | 0.076 | 0.175 |
|  |  | globalEff | 0.194 | 0.163 |
|  |  | maxEigenvalue | 0.600 | 0.227 |
|  |  | medianWeight | 0.142 | 0.046 |
|  |  | modularity | 0.274 | 0.674 |
|  |  | numComponents | 0.766 | 0.766 |
|  |  | numEdges | 0.731 | 0.780 |
|  |  | numModules | 0.476 | 0.014 |
|  |  | numNodes | 0.733 | 0.733 |
|  |  | radius | 0.669 | 0.330 |
|  |  | transitivity | 0.317 | 0.127 |
|  | Offset | assortCoeff |  |  |
|  |  | avgBetweenCentrality | 0.161 | 0.065 |
|  |  | avgClusteringCoeff | 0.799 | 0.537 |
|  |  | avgDegDist | 0.584 | 0.871 |
|  |  | avgEigenvalue | 0.835 | 0.799 |
|  |  | avgLocalEff | 0.756 | 0.881 |
|  |  | avgNeighborDeg | 0.973 | 0.381 |
|  |  | avgParticipationCoeff | 0.022 | 0.830 |
|  |  | avgShortestPaths | 0.334 | 0.139 |
|  |  | avgWeight | 0.814 | 0.570 |
|  |  | density | 0.569 | 0.889 |
|  |  | diameter | 0.469 | 0.031 |
|  |  | globalEff | 0.920 | 0.503 |
|  |  | maxEigenvalue | 0.554 | 0.570 |
|  |  | medianWeight | 0.535 | 0.526 |
|  |  | modularity | 0.250 | 0.041 |
|  |  | numComponents | 0.056 | 0.056 |
|  |  | numEdges | 0.196 | 0.176 |
|  |  | numModules | 0.213 | 0.482 |
|  |  | numNodes | 0.110 | 0.110 |
|  |  | radius | 0.672 | 0.340 |
|  |  | transitivity | 0.924 | 0.513 |
| Identity | Onset | assortCoeff |  |  |
|  |  | avgBetweenCentrality | 0.481 | 0.001 |
|  |  | avgClusteringCoeff | 0.010 | 0.060 |
|  |  | avgDegDist | 0.577 | 0.641 |
|  |  | avgEigenvalue | 0.017 | 0.020 |
|  |  | avgLocalEff | 0.009 | 0.005 |
|  |  | avgNeighborDeg | 0.066 | 0.002 |
|  |  | avgParticipationCoeff | 0.001 | 0.204 |
|  |  | avgShortestPaths | 0.013 | 0.005 |
|  |  | avgWeight | 0.017 | 0.003 |
|  |  | density | 0.549 | 0.661 |
|  |  | diameter | 0.000 | 0.004 |
|  |  | globalEff | 0.028 | 0.006 |
|  |  | maxEigenvalue | 0.211 | 0.005 |
|  |  | medianWeight | 0.008 | 0.001 |
|  |  | modularity | 0.020 | 0.000 |
|  |  | numComponents | 0.032 | 0.032 |
|  |  | numEdges | 0.287 | 0.002 |
|  |  | numModules | 0.033 | 0.031 |
|  |  | numNodes | 0.003 | 0.003 |
|  |  | radius | 0.470 | 0.166 |
|  |  | transitivity | 0.028 | 0.064 |
|  | Offset | assortCoeff |  |  |
|  |  | avgBetweenCentrality | 0.219 | 0.065 |
|  |  | avgClusteringCoeff | 0.053 | 0.023 |
|  |  | avgDegDist | 0.391 | 0.302 |
|  |  | avgEigenvalue | 0.097 | 0.004 |
|  |  | avgLocalEff | 0.034 | 0.008 |
|  |  | avgNeighborDeg | 0.033 | 0.017 |
|  |  | avgParticipationCoeff | 0.025 | 0.920 |
|  |  | avgShortestPaths | 0.078 | 0.072 |
|  |  | avgWeight | 0.046 | 0.013 |
|  |  | density | 0.395 | 0.302 |
|  |  | diameter | 0.044 | 0.042 |
|  |  | globalEff | 0.034 | 0.013 |
|  |  | maxEigenvalue | 0.034 | 0.008 |
|  |  | medianWeight | 0.025 | 0.004 |
|  |  | modularity | 0.192 | 0.018 |
|  |  | numComponents | 0.041 | 0.041 |
|  |  | numEdges | 0.020 | 0.062 |
|  |  | numModules | 0.598 | 0.619 |
|  |  | numNodes | 0.030 | 0.030 |
|  |  | radius | 0.542 | 0.317 |
|  |  | transitivity | 0.077 | 0.023 |

**Table S7.** Classifier results from permutation testing on full networks. Best performance achieved by Logistic Regression classifier on a specific classification task – namely, for a given session, pulse switch type, comparison, and one of three sets of features, correctly classify responses. The leave-one-out cross validation accuracy, the mean and standard deviation of the accuracies of a null distribution built using 1000 permutations of the labels, and the corresponding p-value, or relative position of its accuracy in the null distribution, are all listed. Values in red attained significantly above-chance accuracies. Some tasks did not exceed chance (e.g., Valence on stimulus onset during Stimulus sessions with activity features), and this is indicated by a dashed line to indicate that no permutation testing was conducted. Accuracy on some classification tasks was higher when features were standardized (y = yes, n = no, y/n = same accuracy with or without standardization). Chance is 70% for Valence and 20% for Identity. GT = Graph Theory, Comb = Activity + Graph Theory.

| Session | Pulse Switch | Comparison | Features | Best Possible Accuracy (%) | Permutation score (%<br>mean±s.d.) | p-value | Standardized |
| --- | --- | --- | --- | --- | --- | --- | --- |
| Buffer | Onset | Valence | GT | 70 | - | - | n |
|  |  |  | Activity | 70 | - | - | n |
|  |  |  | Comb. | 70 | - | - | n |
|  |  | Identity | GT | 20 | - | - | y |
|  |  |  | Activity | 17 | - | - | y |
|  |  |  | Comb. | 13 | - | - | y |
|  | Offset | Valence | GT | 73 | 61±7 | 0.074 | y |
|  |  |  | Activity | 70 | - | - | n |
|  |  |  | Comb. | 73 | 58±9 | 0.066 | y |
|  |  | Identity | GT | 33 | 16±8 | 0.021 | y |
|  |  |  | Activity | 13 | - | - | y |
|  |  |  | Comb. | 27 | 11±7 | 0.032 | n |
| Stimulus | Onset | Valence | GT | 70 | - | - | y/n |
|  |  |  | Activity | 67 | - | - | n |
|  |  |  | Comb. | 73 | 59±9 | 0.069 | y |
|  |  | Identity | GT | 40 | 16±8 | 0.004 | y |
|  |  |  | Activity | 17 | - | - | y |
|  |  |  | Comb. | 47 | 16±8 | 0.003 | y |
|  | Offset | Valence | GT | 70 | - | - | n |
|  |  |  | Activity | 70 | - | - | n |
|  |  |  | Comb. | 67 | - | - | y |
|  |  | Identity | GT | 17 | - | - | y |
|  |  |  | Activity | 23 | 15±8 | 0.196 | y |
|  |  |  | Comb. | 33 | 16±8 | 0.038 | y |

**Table S8.** Classifier results from permutation testing on networks with the top 30% of weights. Best performance achieved by Logistic Regression classifier on a specific classification task – namely, for a given session, pulse switch type, comparison, and one of three sets of features, correctly classify responses. The leave-one-out cross validation accuracy, the mean and standard deviation of the accuracies of a null distribution built using 1000 permutations of the labels, and the corresponding p-value, or relative position of its accuracy in the null distribution, are all listed. Values in red attained significantly above-chance accuracies. Some tasks did not exceed chance (e.g., Valence on stimulus onset during Stimulus sessions with activity features), and this is indicated by a dashed line to indicate that no permutation testing was conducted. Accuracy on some classification tasks was higher when features were standardized (y = yes, n = no, y/n = same accuracy with or without standardization). Chance is 70% for Valence and 20% for Identity. GT = Graph Theory, Comb = Activity + Graph Theory.

| Session | Pulse Switch | Comparison | Features | Best Possible Accuracy (%) | Permutation score (%<br>mean±s.d.) | p-value | Standardized |
| --- | --- | --- | --- | --- | --- | --- | --- |
| Buffer | Onset | Valence | GT | 67 | - | - | n |
|  |  |  | Activity | 67 | - | - | y |
|  |  |  | Comb. | 70 | - | - | n |
|  |  | Identity | GT | 13 | - | - | n |
|  |  |  | Activity | 17 | - | - | y |
|  |  |  | Comb. | 23 | 17±8 | 0.265 | y |
|  | Offset | Valence | GT | 57 | - | - | n |
|  |  |  | Activity | 70 | - | - | n |
|  |  |  | Comb. | 60 | - | - | n |
|  |  | Identity | GT | 23 | 15±8 | 0.209 | y |
|  |  |  | Activity | 13 | - | - | y |
|  |  |  | Comb. | 20 | - | - | n |
| Stimulus | Onset | Valence | GT | 73 | 60±8 | 0.065 | y/n |
|  |  |  | Activity | 67 | - | - | n |
|  |  |  | Comb. | 73 | 58±9 | 0.054 | y/n |
|  |  | Identity | **GT** | **47** | **14±8** | **0.001** | **y/n** |
|  |  |  | Activity | 17 | - | - | y |
|  |  |  | **Comb.** | **43** | **14±8** | **0.002** | **n** |
|  | Offset | Valence | GT | 63 | - | - | n |
|  |  |  | Activity | 70 | - | - | n |
|  |  |  | Comb. | 67 | - | - | y |
|  |  | Identity | GT | 27 | 16±8 | 0.115 | y |
|  |  |  | Activity | 23 | 15±8 | 0.196 | y |
|  |  |  | **Comb.** | **37** | **16±8** | **0.022** | **y** |

**Table S9.**
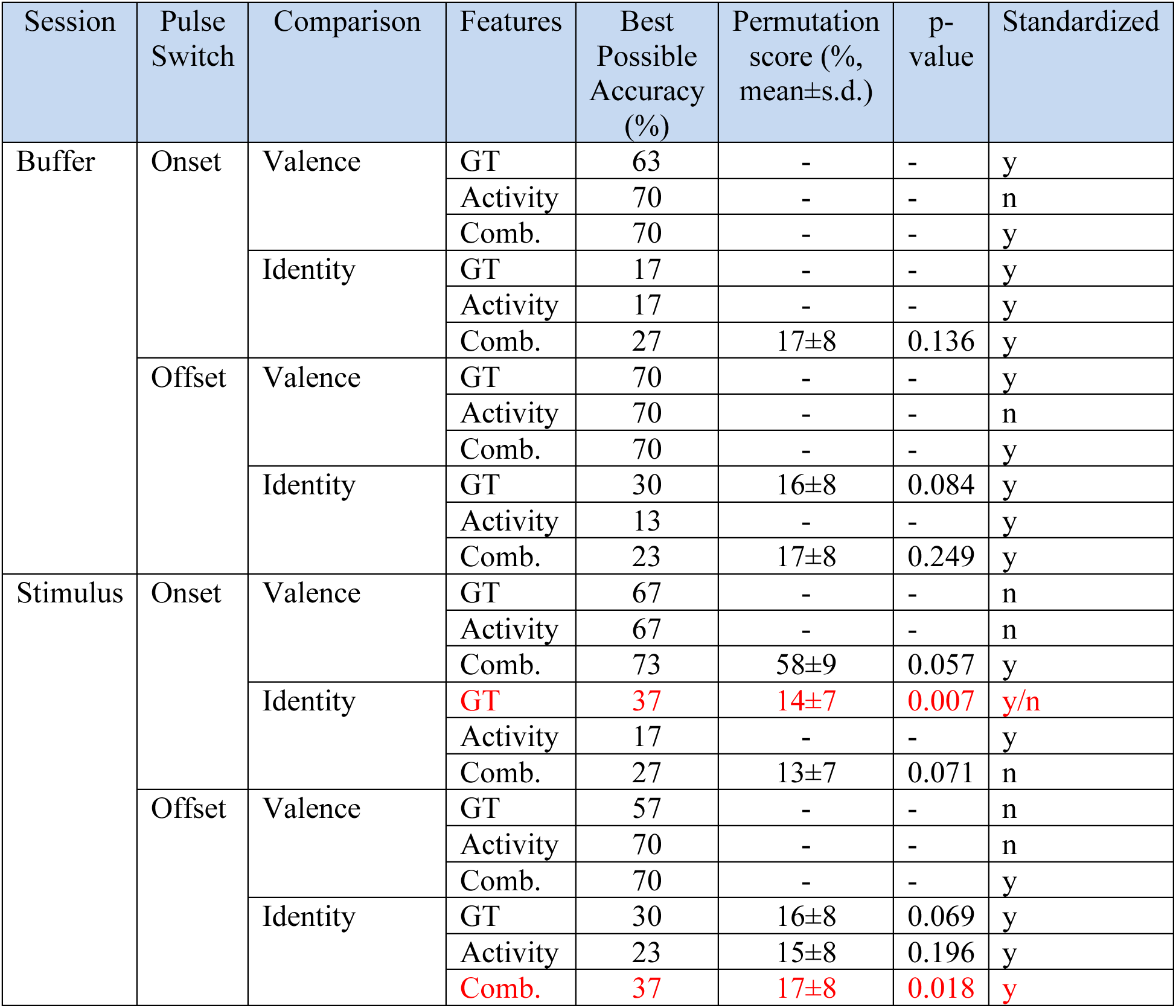
Classifier results from permutation testing on networks with the top 20% of weights. Best performance achieved by Logistic Regression classifier on a specific classification task – namely, for a given session, pulse switch type, comparison, and one of three sets of features, correctly classify responses. The leave-one-out cross validation accuracy, the mean and standard deviation of the accuracies of a null distribution built using 1000 permutations of the labels, and the corresponding p-value, or relative position of its accuracy in the null distribution, are all listed. Values in red attained significantly above-chance accuracies. Some tasks did not exceed chance (e.g., Valence on stimulus onset during Stimulus sessions with activity features), and this is indicated by a dashed line to indicate that no permutation testing was conducted. Accuracy on some classification tasks was higher when features were standardized (y = yes, n = no, y/n = same accuracy with or without standardization). Chance is 70% for Valence and 20% for Identity. GT = Graph Theory, Comb = Activity + Graph Theory.

| Session | Pulse Switch | Comparison | Features | Best Possible Accuracy (%) | Permutation score (%<br>mean±s.d.) | p-value | Standardized |
| --- | --- | --- | --- | --- | --- | --- | --- |
| Buffer | Onset | Valence | GT | 63 | - | - | y |
|  |  |  | Activity | 70 | - | - | n |
|  |  |  | Comb. | 70 | - | - | y |
|  |  | Identity | GT | 17 | - | - | y |
|  |  |  | Activity | 17 | - | - | y |
|  |  |  | Comb. | 27 | 17±8 | 0.136 | y |
|  | Offset | Valence | GT | 70 | - | - | y |
|  |  |  | Activity | 70 | - | - | n |
|  |  |  | Comb. | 70 | - | - | y |
|  |  | Identity | GT | 30 | 16±8 | 0.084 | y |
|  |  |  | Activity | 13 | - | - | y |
|  |  |  | Comb. | 23 | 17±8 | 0.249 | y |
| Stimulus | Onset | Valence | GT | 67 | - | - | n |
|  |  |  | Activity | 67 | - | - | n |
|  |  |  | Comb. | 73 | 58±9 | 0.057 | y |
|  |  | Identity | GT | 37 | 14±7 | 0.007 | y/n |
|  |  |  | Activity | 17 | - | - | y |
|  |  |  | Comb. | 27 | 13±7 | 0.071 | n |
|  | Offset | Valence | GT | 57 | - | - | n |
|  |  |  | Activity | 70 | - | - | n |
|  |  |  | Comb. | 70 | - | - | y |
|  |  | Identity | GT | 30 | 16±8 | 0.069 | y |
|  |  |  | Activity | 23 | 15±8 | 0.196 | y |
|  |  |  | Comb. | 37 | 17±8 | 0.018 | y |

**Table S10.** Classifier results from permutation testing on networks with the top 10% of weights. Best performance achieved by Logistic Regression classifier on a specific classification task – namely, for a given session, pulse switch type, comparison, and one of three sets of features, correctly classify responses. The leave-one-out cross validation accuracy, the mean and standard deviation of the accuracies of a null distribution built using 1000 permutations of the labels, and the corresponding p-value, or relative position of its accuracy in the null distribution, are all listed. Values in red attained significantly above-chance accuracies. Some tasks did not exceed chance (e.g., Valence on stimulus onset during Stimulus sessions with activity features), and this is indicated by a dashed line to indicate that no permutation testing was conducted. Accuracy on some classification tasks was higher when features were standardized (y = yes, n = no, y/n = same accuracy with or without standardization). Chance is 70% for Valence and 20% for Identity. GT = Graph Theory, Comb = Activity + Graph Theory.

| Session | Pulse Switch | Comparison | Features | Best Possible Accuracy (%) | Permutation score (%<br>mean±s.d.) | p-value | Standardized |
| --- | --- | --- | --- | --- | --- | --- | --- |
| Buffer | Onset | Valence | GT | 67 | - | - | n |
|  |  |  | Activity | 70 | - | - | n |
|  |  |  | Comb. | 67 | - | - | n |
|  |  | Identity | GT | 23 | 14±8 | 0.148 | n |
|  |  |  | Activity | 17 | - | - | y |
|  |  |  | Comb. | 27 | 17±8 | 0.155 | y |
|  | Offset | Valence | GT | 57 | - | - | y |
|  |  |  | Activity | 70 | - | - | n |
|  |  |  | Comb. | 57 | - | - | n |
|  |  | Identity | GT | 13 | - | - | n |
|  |  |  | Activity | 13 | - | - | y |
|  |  |  | Comb. | 13 | - | - | y |
| Stimulus | Onset | Valence | GT | 60 | - | - | n |
|  |  |  | Activity | 67 | - | - | n |
|  |  |  | Comb. | 63 | - | - | y |
|  |  | Identity | GT | 23 | 16±8 | 0.215 | y |
|  |  |  | Activity | 17 | - | - | y |
|  |  |  | Comb. | 30 | 16±7 | 0.069 | y |
|  | Offset | Valence | GT | 70 | - | - | n |
|  |  |  | Activity | 60 | - | - | y |
|  |  |  | Comb. | 60 | - | - | y/n |
|  |  | Identity | GT | 20 | - | - | y |
|  |  |  | Activity | 23 | 15±8 | 0.196 | y |
|  |  |  | Comb. | 23 | 16±8 | 0.215 | y |

**Table S11.** Classifier results from permutation testing on networks with positively correlated neurons. Best performance achieved by Logistic Regression classifier on a specific classification task – namely, for a given session, pulse switch type, comparison, and one of three sets of features, correctly classify responses. The leave-one-out cross validation accuracy, the mean and standard deviation of the accuracies of a null distribution built using 1000 permutations of the labels, and the corresponding p-value, or relative position of its accuracy in the null distribution, are all listed. Values in red attained significantly above-chance accuracies. Some tasks did not exceed chance (e.g., Valence on stimulus onset during Stimulus sessions with activity features), and this is indicated by a dashed line to indicate that no permutation testing was conducted. Accuracy on some classification tasks was higher when features were standardized (y = yes, n = no, y/n = same accuracy with or without standardization). Chance is 70% for Valence and 20% for Identity. GT = Graph Theory, Comb = Activity + Graph Theory.

| Session | Pulse Switch | Comparison | Features | Best Possible Accuracy (%) | Permutation score (%<br>mean±s.d.) | p-value | Standardized |
| --- | --- | --- | --- | --- | --- | --- | --- |
| Buffer | Onset | Valence | GT | 63 | - | - | n |
|  |  |  | Activity | 70 | - | - | n |
|  |  |  | Comb. | 63 | - | - | n |
|  |  | Identity | GT | 17 | - | - | y |
|  |  |  | Activity | 17 | - | - | y |
|  |  |  | Comb. | 23 | 16±7 | 0.235 | y |
|  | Offset | Valence | GT | 67 | - | - | y/n |
|  |  |  | Activity | 70 | - | - | n |
|  |  |  | Comb. | 63 | - | - | y/n |
|  |  | Identity | GT | 23 | 16±8 | 0.236 | y |
|  |  |  | Activity | 13 | - | - | y |
|  |  |  | Comb. | 20 | - | - | n |
| Stimulus | Onset | Valence | GT | 67 | - | - | y/n |
|  |  |  | Activity | 67 | - | - | n |
|  |  |  | Comb. | 60 | - | - | y/n |
|  |  | Identity | **GT** | **40** | **14±8** | **0.003** | **n** |
|  |  |  | Activity | 17 | - | - | y |
|  |  |  | **Comb.** | **40** | **15±8** | **0.004** | **n** |
|  | Offset | Valence | GT | 73 | 59±8 | 0.064 | y |
|  |  |  | Activity | 70 | - | - | n |
|  |  |  | Comb. | 70 | - | - | n |
|  |  | Identity | GT | 27 | 16±7 | 0.102 | y |
|  |  |  | Activity | 23 | 15±8 | 0.196 | y |
|  |  |  | Comb. | 30 | 16±7 | 0.061 | y |

**Table S12.** Classifier results from permutation testing on networks with negatively correlated neurons. Best performance achieved by Logistic Regression classifier on a specific classification task – namely, for a given session, pulse switch type, comparison, and one of three sets of features, correctly classify responses. The leave-one-out cross validation accuracy, the mean and standard deviation of the accuracies of a null distribution built using 1000 permutations of the labels, and the corresponding p-value, or relative position of its accuracy in the null distribution, are all listed. Values in red attained significantly above-chance accuracies. Some tasks did not exceed chance (e.g., Valence on stimulus onset during Stimulus sessions with activity features), and this is indicated by a dashed line to indicate that no permutation testing was conducted. Accuracy on some classification tasks was higher when features were standardized (y = yes, n = no, y/n = same accuracy with or without standardization). Chance is 70% for Valence and 20% for Identity. GT = Graph Theory, Comb = Activity + Graph Theory.

| Session | Pulse Switch | Comparison | Features | Best Possible Accuracy (%) | Permutation score (%<br>mean±s.d.) | p-value | Standardized |
| --- | --- | --- | --- | --- | --- | --- | --- |
| Buffer | Onset | Valence | GT | 63 | - | - | y |
|  |  |  | Activity | 70 | - | - | n |
|  |  |  | Comb. | 63 | - | - | y |
|  |  | Identity | GT | 20 | - | - | y |
|  |  |  | Activity | 17 | - | - | y |
|  |  |  | Comb. | 20 | - | - | y |
|  | Offset | Valence | GT | 63 | - | - | n |
|  |  |  | Activity | 70 | - | - | n |
|  |  |  | Comb. | 63 | - | - | n |
|  |  | Identity | GT | 23 | 16±7 | 0.234 | y |
|  |  |  | Activity | 13 | - | - | y |
|  |  |  | Comb. | 20 | - | - | n |
| Stimulus | Onset | Valence | GT | 77 | 60±8 | 0.040 | y |
|  |  |  | Activity | 67 | - | - | n |
|  |  |  | Comb. | 73 | 58±9 | 0.071 | y/n |
|  |  | Identity | GT | 33 | 16±8 | 0.034 | y |
|  |  |  | Activity | 17 | - | - | y |
|  |  |  | Comb. | 30 | 14±8 | 0.051 | y/n |
|  | Offset | Valence | GT | 67 | - | - | n |
|  |  |  | Activity | 70 | - | - | n |
|  |  |  | Comb. | 70 | - | - | y |
|  |  | Identity | GT | 20 | - | - | y |
|  |  |  | Activity | 23 | 15±8 | 0.196 | y |
|  |  |  | Comb. | 33 | 16±7 | 0.031 | y |

